# Platelet integrin αIIbβ3 plays a key role in venous thrombogenesis in a mouse model

**DOI:** 10.1101/2024.07.11.602533

**Authors:** Brian D. Adair, Conroy O. Field, José L. Alonso, Jian-Ping Xiong, Shi-Xian Deng, Hyun Sook Ahn, Eivgeni Mashin, Clary B. Clish, Johannes van Agthoven, Mark Yeager, Youzhong Guo, David A Tess, Donald W Landry, Mortimer Poncz, M. Amin Arnaout

## Abstract

Venous thrombosis (VT) is a common vascular disease associated with reduced survival and a high recurrence rate. Previous studies have shown that the accumulation of platelets and neutrophils at sites of endothelial cell activation is a primary event in VT, but a role for platelet αIIbβ3 in the initiation of venous thrombosis has not been established. This task has been complicated by the increased bleeding linked to partial agonism of current αIIbβ3 inhibitory drugs such as tirofiban (Aggrastat^®^). Here, we show that m-tirofiban, an engineered version of tirofiban, is not a partial agonist of αIIbβ3. This is based on its cryo-EM structure in complex with human full-length αIIbβ3 and its inability to increase expression of an activation-sensitive epitope on platelet αIIbβ3. m-tirofiban abolished agonist-induced platelet aggregation *ex vivo* at concentrations that preserved clot retraction and markedly suppressed the accumulation of platelets, neutrophils, and fibrin on thrombin-activated endothelium in real-time using intravital microscopy in a mouse model of venous thrombogenesis. Unlike tirofiban, however, m-tirofiban did not increase bleeding at the thrombosis-inhibitory dose. These findings establish a key role for αIIbβ3 in the initiation of VT, provide a guiding principle for designing potentially safer inhibitors for other integrins, and suggest that pure antagonists of αIIbβ3 like m-tirofiban merit further consideration as potential thromboprophylaxis agents in patients at high-risk for VT and hemorrhage.

## Introduction

Venous thrombosis (VT) is the third most common vascular disease in humans, with an annual incidence of 1-2 per 1,000 population, rising to above 1 per 100 hospitalized patients ^1^ and up to 20% of hospitalized COVID-19 patients ^2^. VT is associated with increased short- and long-term mortality risk and a high recurrence rate, with estimated healthcare costs of 7-10 billion dollars in the USA^3^. Predisposing factors, in addition to older age and COVID-19, include major surgery, active cancer, prolonged immobilization, obesity, and atherosclerosis ^4^. Anticoagulant therapy is the cornerstone of treatment but increases bleeding risk, especially in post-surgical patients or following major trauma ^5^, underscoring the need for preventative measures in high-risk patients.

VT is a thromboinflammatory disease. Prolonged blood stasis converts intact venous endothelium, primarily in the sinus of the venous valves of large veins such as the femoral vein ^6^ into a procoagulant proinflammatory phenotype recruiting platelets and innate immune cells, which release procoagulant factors such as tissue factor and prothrombotic neutrophil extracellular traps (NETs), contributing to the formation and growth of venous thrombi ^5,7^. Several studies have suggested that monotherapy with the antiplatelet drug aspirin is partially effective in preventing VT ^8–10^, with more potent inhibitors of platelet protease-activated receptor-1 (PAR-1)- and ADP receptor (P2Y_12_), further lowering the VT risk by ∼30% relative to aspirin alone ^11^. PAR-1 and P2Y_12_ receptors are known to activate platelet integrin αIIbβ3, a receptor that plays a central role in arterial thrombosis ^12^, suggesting that αIIbβ3 may also contribute to the pathophysiology of VT. However, the increased bleeding associated with the use of direct αIIbβ3 inhibitory drugs, which is linked to inadvertent drug-induced activating conformational changes in the receptor ^13^, has hampered investigations into a potential pathogenic role of αIIbβ3 in VT that could be targeted therapeutically.

In this study, we used m-tirofiban, an engineered version of the specific and direct αIIbβ3 inhibitor drug tirofiban, to evaluate the potential role of αIIbβ3 in the initiation of VT. In contrast with tirofiban, m-tirofiban does not induce the activating conformational changes in the receptor as revealed in its cryo-EM structure in complex with human full-length αIIbβ3 and by the lack of increased expression of the activation-sensitive LIBS (ligand-induced binding site) epitope AP5 ^14^ on human platelets. m-tirofiban inhibited ADP-induced human platelet aggregation *ex vivo* at a half-maximal aggregation inhibitory concentration (*IC_50 agg_*) that was 1,020 times lower than the half-maximal clot retraction inhibitory concentration (*IC_50 clot_*), compared to the very narrow *IC_50 clot_/IC_50 agg_* ratio of 27.0 for tirofiban. Pretreatment of wild-type C57BL/6J mice with a loading dose of m-tirofiban inhibited venous thrombogenesis in a mouse model but did not increase bleeding, unlike tirofiban. These findings establish a key role for αIIbβ3 in the initiation of VT and suggest that m-tirofiban may prove useful in thromboprophylaxis of VT-prone patients where anticoagulants are contraindicated.

## Results

### Synthesis of m-tirofiban

m-tirofiban ((S)-m-tirofiban) is a modification of the tirofiban structure in which the sulfonamide-butane moiety of tirofiban is replaced with benzoxazole (**Supplementary Fig. 1a**). Introduction of the benzoxazole moiety is designed to mimic a key tryptophan residue in fibronectin-10-derived hFN10 and Hr10 ligands previously shown responsible for preventing the activating conformational changes in hFN10- or Hr10-bound β3 integrins ^15,16^. m-tirofiban was prepared by the synthetic route described under “Methods.” The structure of the product was confirmed via proton and carbon NMR spectroscopy and mass spectrometry. LC/MS showed a single peak with a retention time of 7.5 min and m/z 466 (M+1); with no contamination by hydrolysis product(s) (**Supplementary Fig.1b-d**). A 10 mM stock solution in anhydrous DMSO was stored in aliquots at -80° C.

### Cryo-EM structures of full-length human αIIbβ3 in complex with m-tirofiban or tirofiban

Previous studies determined the structures of αIIbβ3 in lipid bilayer nanodiscs as the membrane mimetic ^17,18^. These cryo-EM maps revealed important conformational states, but the resolution was too low to visualize secondary structural elements. In contrast, we used the NCMNP7b polymer ^19^ to isolate patches of native platelet membrane, obviating the need for detergent solubilization and reconstitution in nanodiscs ^20^. The cryo-EM map of m-tirofiban-bound to full-length αIIbβ3 had an overall resolution of 4.1 Å, with a maximal resolution of 3.4 Å for the head domain (comprising the αIIb propeller domain and the β3 βA domain) (Fig. 1a, Supplemental Table 1, Supplemental Fig. 2). m-tirofiban-bound αIIbβ3 recapitulates the canonical bent inactive conformation as that of the unliganded full-length αIIbβ3 ^20^ (**Supplementary Fig. 3**). m-tirofiban engages the Arg-Gly-Asp binding site in the integrin head with the nitrogen of the Arg-mimetic piperidine H-bonding to the propeller’s Asp^224^ and one of its Asp-mimetic carboxyl oxygens coordinating the Mg^2+^ at the *m*etal-*i*on-*d*ependent-*a*dhesion-*s*ite (MIDAS) of the βA domain (**Fig. 1b**). The distances of m-tirofiban’s benzoxazole to the side chains of Tyr^122^ and Arg^214^ of βA, can be explained by a π- π and a cation- π interactions, respectively, as these distances are consistent with those in published reports ^21,22^. These interactions prevent the activating inward movement of the α1 helix towards the MIDAS motif while maintaining the ADMIDAS Ca^2+^-mediated link between the α1 helix and the strand F- α7 loop, thus freezing the βA domain in the inactive conformation and hence freezing the integrin in the inactive state ^23^.

**Figure 1.**
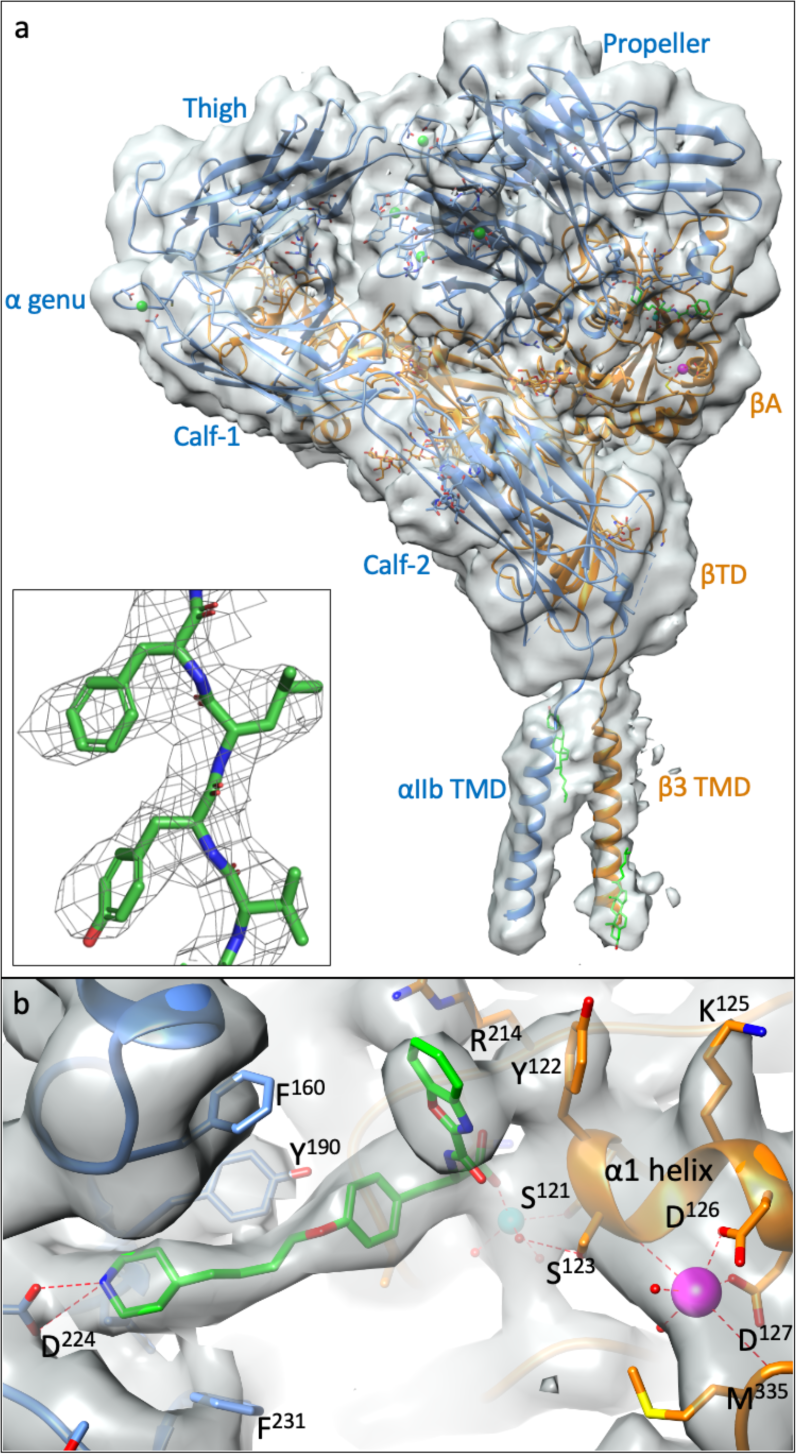
Cryo-EM structure of the full-length αIIbβ3/m-tirofiban complex. **a** The unsharpened cryo-EM map at 4.1 Å, displayed with Chimera, is shown with ribbon diagrams for the αIIb chain (light blue), β3 chain (orange), and m-tirofiban (green). All 12 subdomains of the ectodomain are resolved, and the αIIb and β3 TM domains (TMD) are visualized. The additional densities in the TM map have been modeled as cholesterol molecules (carbons in green, oxygens in red). The four calcium ions at the base of the propeller and the one at the α-genu are shown as green spheres. The four extracellular αIIb and two (βA and βTD) of the 8 β3 domains are labeled. Inset, cryo-EM map (displayed with PyMOL at a contour level of 14) showing a region of secondary structure (residues V^328^-F^331^). **b** Closeup view of the EM density of m-tirofiban. To improve the view of the side chains, the map has been sharpened with an applied B-factor of 150 Å^2^. The m-tirofiban-integrin contacts are described in the text. The metal ions at MIDAS and ADMIDAS are shown as cyan and magenta spheres, respectively.

In contrast, the cryo-EM structure of the tirofiban-bound αIIbβ3 shows large conformational changes in the integrin headpiece (comprising the integrin head, αIIb’s thigh domain, and β3’s hybrid, PSI, and EGF1 domains), which adopts the open (hybrid-swingout) conformation (**Fig. 2a**, Supplementary Table 1 and Supplementary Fig. 4), also seen in the αIIbβ3 cryo-EM structure in complex with another partial agonist drug eptifibatide (Integrilin^®^) ^20^. The αIIb’s thigh domain is partially visualized, but the TM domains are not, presumably due to conformational flexibility. During the initial particle selection, one of the 2D classes, containing 19,994 particles (out of a total of 975,560 selected), displayed a view of the bent conformation (**Supplementary Fig. 4b**), indicating that this minor subpopulation maintains the bent conformation despite prolonged (>24 hrs) incubation with tirofiban. In the unbent conformation, tirofiban occupies the Arg-Gly-Asp binding site with its sulfonamide oxygens contacting βA’s Arg^214^ epsilon nitrogen (NE) and Tyr^166^ and αIIb propeller’s Tyr^190^ (**Fig. 2b**). The swingout of the hybrid domain underneath of βA is allowed by the unrestricted inward movement of the α1 helix towards the MIDAS motif that breaks the inactivating ADMIDAS Ca^2+^-mediated link between the α1 helix and the strand F- α7 loop (**Fig. 2c**). These drug-induced conformational changes have been linked to adverse outcomes in treated patients, limiting the use of partial agonist drugs to high-risk settings for arterial thrombosis ^13^.

**Figure 2.**
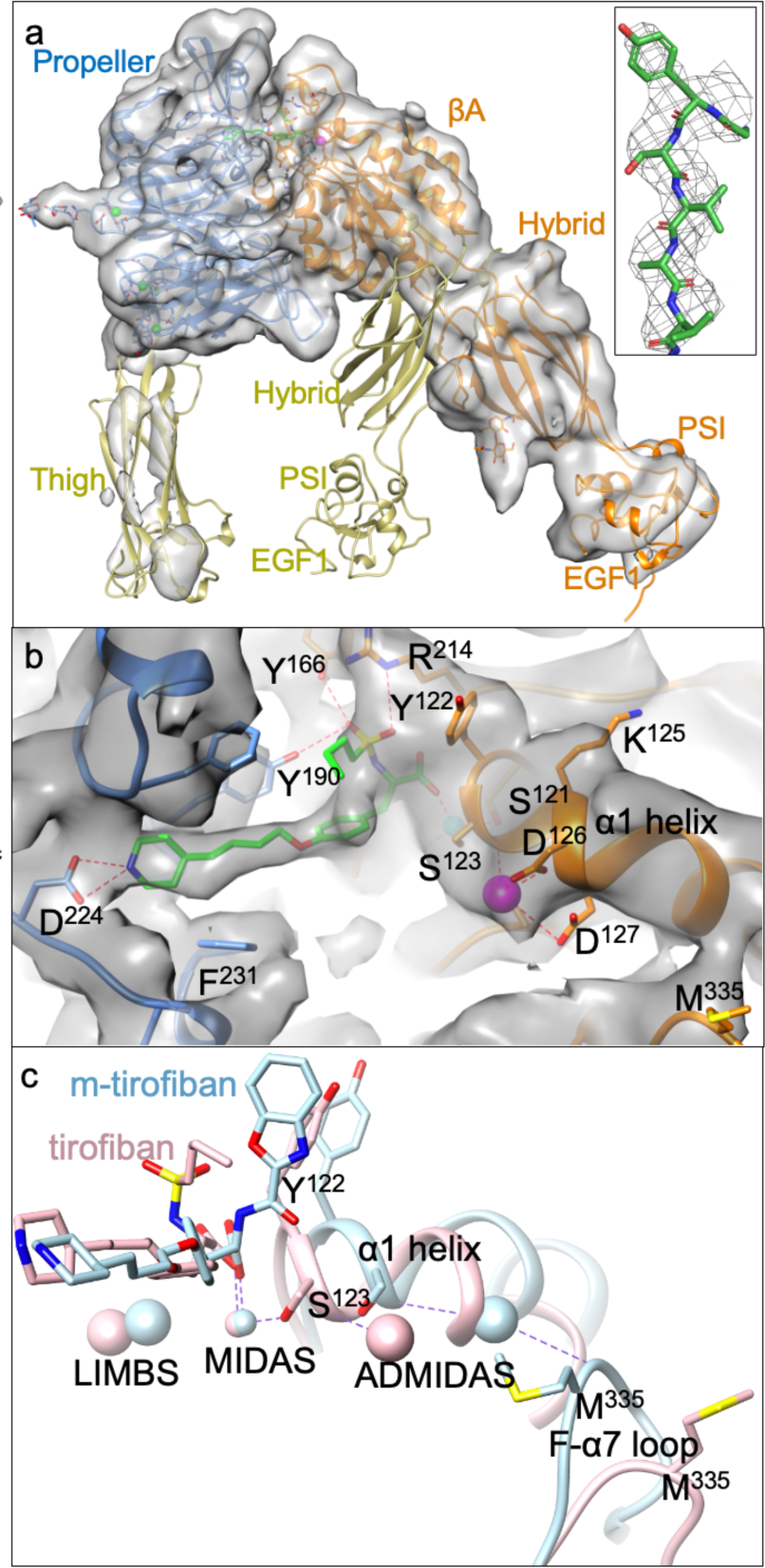
Cryo-EM structure of the full-length αIIbβ3/tirofiban complex. **a** The unsharpened cryo-EM map of tirofiban-bound αIIbβ3 is displayed with Chimera with αIIb in light blue, β3 in orange, and tirofiban in green. The map, at the overall resolution of 3.9 Å, shows only a density for the integrin αIIb propeller, βA-, hybrid- and PSI domains with a weak density for the αIIb thigh domain and β3 EGF1 but no densities for the leg (αIIb’s Calf1, 2 and β3’s EGF2-4 and βTD)- and TM domains. The four calcium ions at the base of the propeller are shown as green spheres. The αIIb propeller is superposed onto that of the full-length unliganded cryo-EM structure (8t2v.pdb)(khaki) to show the tirofiban-induced swingout of the hybrid domain. Inset, cryo-EM map (displayed with PyMOL at a contour level of 14) showing a region of secondary structure (residues Y^237^-V^241^). **b** Closeup view of the EM density for bound tirofiban in the same orientation as Fig. 1b. To improve the view of the side chains, the map has been sharpened with an applied B-factor of 150 Å^2^. The m-tirofiban-integrin contacts are described in the text. The MIDAS Mg^2+^ and ADMIDAS Ca^2+^ ions are shown as cyan and magenta spheres, respectively. **c** Ribbon tracings of the superposed βA domains of tirofiban/αIIbβ3 and m-tirofiban/αIIbβ3 complexes (in rose and light blue, respectively, with metal ions in the respective colors) showing the key activating tertiary changes in backbone and ADMIDAS metal ion coordination induced by bound tirofiban. The α1 helix and stand F-α7 loop are shown as thick and thinner tubes. Oxygens are red, nitrogens are blue, and sulfurs are yellow.

### Inhibitor binding to platelet-free human plasma

The introduction of the hydrophobic benzoxazole moiety into drugs such as tafamidis ^24^ is known to significantly increase their binding to albumin. We determined the unbound fraction ( *f_u,p_*) of m-tirofiban to platelet-free human plasma using rapid equilibrium dialysis and mass spectrometry, in which tirofiban was used as an internal control (see “Methods”). The *f_u,p_* of m-tirofiban was 0.95±0.21% (mean+ sd, n=3) compared with 41.27±3.03 % for tirofiban (**Supplementary Table 2**), the latter value in good agreement with the 36% *f_u,p_* measured in human blood using ^14^C-labeled tirofiban ^25^. Dissociation constants of tirofiban and m-tirofiban to albumin (*K_d,A_*) of 448.0 µM and 6.11 µM were estimated from the respective mean *f_u,p_* values using a set of ordinary differential equations (ODEs) (Eq1-5 in “Methods” and **Supplementary Fig. 5**).

### Dissociation constants of αIIbβ3 inhibitors

The *IC_50_* of Alexa647-conjugated human fibrinogen (Alexa647-FB) binding to washed human platelets activated with 20 µM ADP in the presence of increasing concentrations of inhibitors was determined in Tyrode’s buffer containing 1 mM each of Ca^2+^ and Mg^2+^ and 0.02% bovine serum albumin (BSA). This analysis yielded *IC_50_*s of 138±15.3 nM for m-tirofiban, 9.95±1.0 nM for tirofiban (consistent with a published report ^26^), and 174.9±12.1 nM for the pure αIIbβ3 antagonist Hr10 ^16^ (mean ± s.e., n=3 donors) (**Fig. 3a**). Dissociation constants (*K_d_*) of the inhibitors to ADP-activated platelet αIIbβ3 were derived from another set of ODEs (Eq6-15 in “Methods” and **Supplementary Fig. 5**), yielding *K_d_* values of 23.0 nM (geometric mean, coefficient of variance of 38%) for m-tirofiban, 1.44 nM (geometric mean, coefficient of variance of 15%) for tirofiban, and 39.7 nM (geometric mean, coefficient of variance of 23%) for Hr10. The derived *K_d_* for tirofiban to active αIIbβ3 agrees with published reports ^26,27^. Our previous study showed that m-tirofiban and Hr10 bind equally to inactive and active αIIbβ3 ^16^.

**Figure 3.**
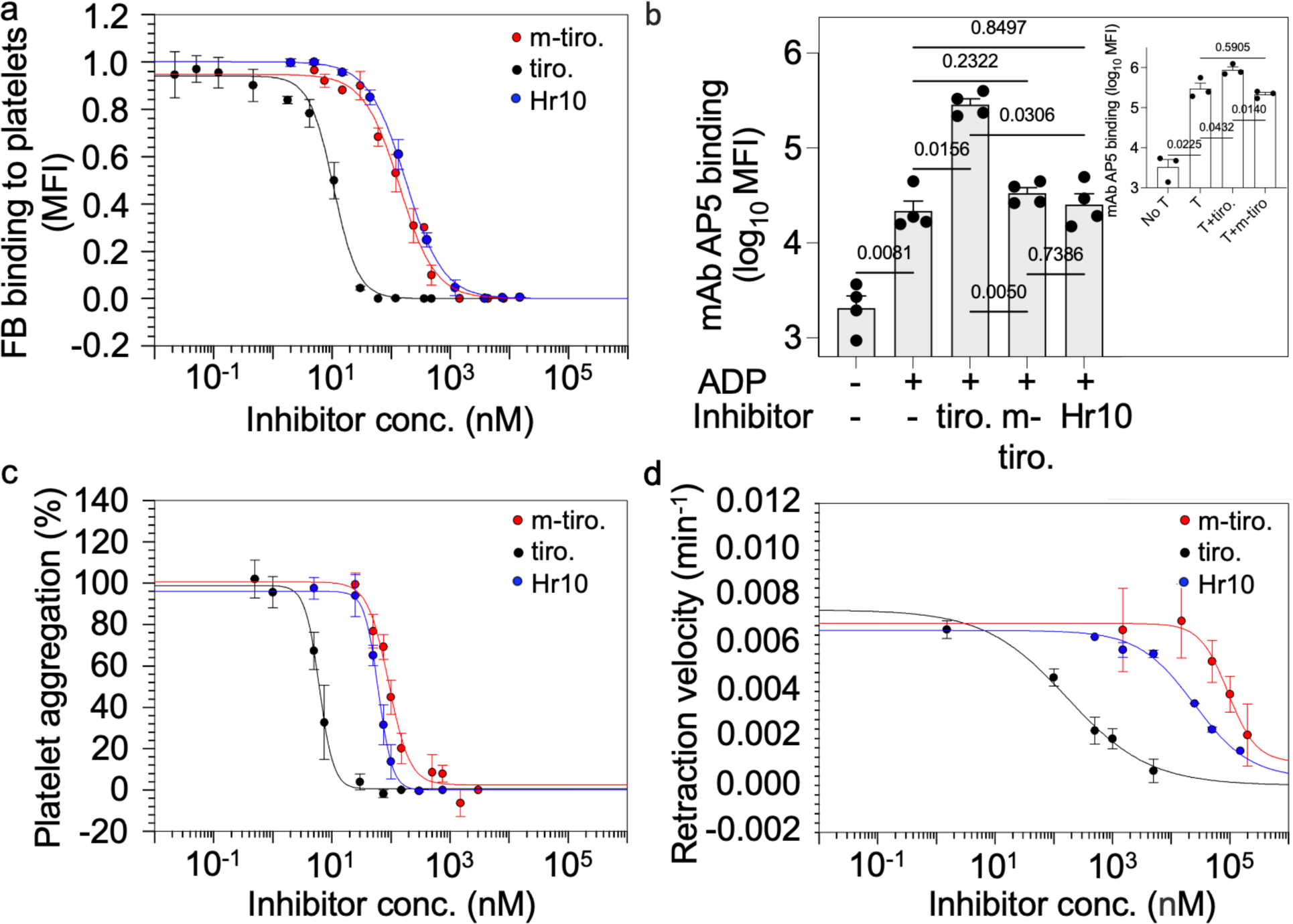
Inhibitor binding properties and effect on αIIbβ3 conformation and function. **a** Dose-response curves (points indicate the mean ± s.e., n = 3 different donors) showing the binding of Alexa647-FB bound to washed ADP-treated human platelets in the presence of increasing concentrations of unlabeled m-tirofiban (m-tiro.), tirofiban (tiro.), or Hr10 and 0.02% BSA. The MFI values from the three FACS analyses were normalized individually before averaging. **b** Bar charts (mean + s.e., n=4 donors) of the expression of LIBS AP5 epitope on human resting platelets and after platelet activation with ADP, alone or in the presence tirofiban (0.17μM), m-tirofiban (16 μM), or Hr10 (7.6 μM) (used at 696x, 118x, and 191x the respective *K_d_* value to active αIIbβ3). AP5 binding is shown as log_10_ mean channel fluorescence (MFI). Inset bar charts (mean + s.e., n=3 donors) showing AP5 binding (log_10_ MFI) to platelets in the absence or presence of thrombin (T) alone or combined with tirofiban (0.17μM) or m-tirofiban (16 μM). Indicated p values are determined by one-way ANOVA with Tukey’s multiple comparison test. **c** Dose-response curves (points indicate the mean ± s.e., 4 donors used for tirofiban and Hr10 and 5 for m-tirofiban) showing the effects of the inhibitors on ADP-induced platelet aggregation in 1:1 diluted whole blood. Points for the integrated impedance from the three experiments were individually normalized prior to averaging and are displayed with least-squares fits to the mean values. **d** Dose-response curves (points indicate the mean ± s.e., n = 3 donors) showing the effects of increasing concentrations of inhibitors on clot retraction velocity.

### Effects of inhibitors on ADP-induced AP5 epitope exposure of αIIbβ3 in human platelets

The tirofiban-induced conformational changes revealed in the tirofiban/αIIbβ3 cryo-EM structure (**Fig. 2a**) expose neoepitopes recognized by LIBS monoclonal antibodies (mAbs) such as AP5 ^14^. As expected, 0.17μM tirofiban, a concentration 118x its *K_d_* value to active αIIbβ3, induced a large increase in AP5 epitope expression above that seen with ADP or thrombin alone (**Fig. 3b**). In contrast, there was no such increase above that induced by ADP alone in the presence of m-tirofiban at 16 μM or Hr10 at 7.6μM, concentrations that are 696x and 191x their respective *K_d_* values to active platelet αIIbβ3 (**Fig. 3b**).

### ADP-induced platelet aggregation

This was performed by preincubating whole blood with an equal volume of saline containing different concentrations of each inhibitor for 5 min at 37°C without stirring, followed by stirring and the addition of ADP at 20 μM, and then measuring platelet aggregation. Tirofiban inhibited ADP-induced platelet aggregation with an *IC_50 agg_* of 6.0 ± 0.58 nM (mean ± se, n=4)(Fig. 3c). Using this value and the tirofiban *K_d_* for pretreated (inactive) αIIbβ3 of 15 nM ^28^, a third set of ODEs (Eq 16-22 in “Methods” and **Supplementary Fig. 5**) estimates that at 26.5 nM (38.0 nM in undiluted whole blood) tirofiban achieves 50% occupancy of αIIbβ3 (T) (*OCC_L:T_*) (i.e., the concentration that inhibits ADP-induced platelet aggregation by 100% ^29^). The 26.5 nM concentration is in excellent agreement with the experimental data (**Fig. 3c**). For an m-tirofiban *IC_50 agg_* of 91.7 ± 9.6 nM (mean ± se, n=5 donors) (**Fig. 3c**), the corresponding 50% *OCC_L:T_* is 688 nM (∼2-fold lower than the experimental value, **Fig. 3c**), and 1.35 μM in undiluted whole blood. For Hr10, with an *IC_50 agg_* of 61.5± 5.1 nM (mean ± se, n=4) (**Fig. 3c**), the 50% *OCC_L:T_* is 45.3 nM (∼4-fold lower than the experimental value, **Fig.3c**), and 51.0 nM in undiluted whole blood. The slightly higher 50% *OCC_L:T_* for Hr10 obtained experimentally (**Fig.3c**) may be due to a relatively slower on-rate for the 12 kDa peptide during the 5-min preincubation period.

### Effect of inhibitors on clot retraction

m-tirofiban inhibited thrombin-induced clot retraction with an *IC_50 clot_* of 93.5 ± 37.5 μM (mean ± se, n=3 donors), compared with 0.162 ± 0.061 μM for tirofiban and 24.8 ± 10.2 μM for Hr10. (**Fig. 3d**). The *IC_50 clot_* /*IC_50 agg_* ratios for m-tirofiban and Hr10 are 1,020 and 403, respectively, the difference likely explained by the albumin binding of m-tirofiban. This ratio is only 27.0 for tirofiban. The need for higher concentrations of all antagonists to block clot retraction vs. aggregation is due to the polyvalent and multisite binding of fibrin fibrils to αIIbβ3 vs the monovalent interaction of FB with the receptor ^30^.

### Thrombin-induced femoral vein thrombosis in C57BL/6J mice

We used a mouse model of venous thrombogenesis under blood flow ^31,32^ to evaluate the potential role of αIIbβ3 in platelet and neutrophil accumulation at sites of endothelial cell activation, a primary event in venous thrombogenesis ^7^. In this model, the intact femoral vein is exposed, the vessel cleared of connective tissue, and thrombin is directly applied to the external vessel wall, followed immediately by quantifying the accumulation of labeled platelets, neutrophils, and fibrin at the activated endothelium in real-time using intravital microscopy (Fig. 4). This model avoids the transluminal injury in the acute ferric chloride VT model and does not impede the effect of systemically administered drugs, otherwise unavoidable in the inferior vena cava ligation/stenosis/stasis models ^33^. It is also well suited to evaluate compounds as potential thromboprophylaxis agents.

**Figure 4.**
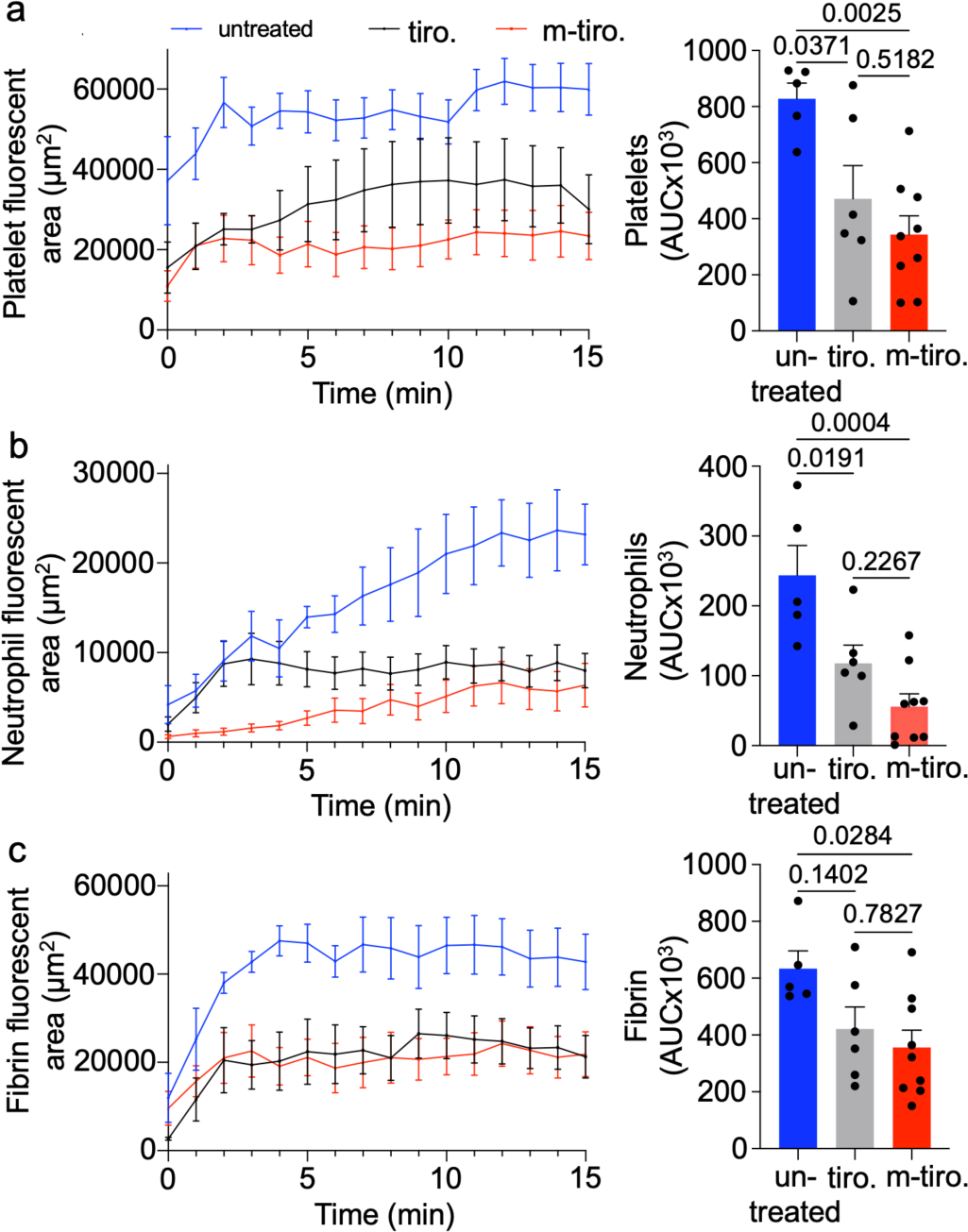
M-tirofiban inhibits femoral vein thrombosis in C57BL/6J mice. Left panels. Graphs showing the kinetics of labeled platelet (a), neutrophil (b), and fibrin (c) accumulation in thrombi in wild-type mice receiving buffer (untreated, blue tracings), tirofiban (0.025μg/gm, gray tracings) or m-tirofiban (0.3μg/gm, red tracings) parenterally five minutes before topical application of thrombin to the exposed intact femoral vein. Tracings on the left display the mean and standard error in fluorescence accumulation. For clarity, only points at 1-minute intervals are displayed. Platelets have already begun accumulating on the activated endothelium in untreated (but not treated) animals (4a) at time zero when video recording usually begins one minute after thrombin application. The right panels show points depicting the integrated area of the full data set for each animal. Bar charts display the mean and s.e. of each group. Each dataset passed the normality test. Indicated p values are determined by one-way ANOVA with Tukey’s test for multiple comparisons.

We compared the thrombin-induced accumulation of platelets, neutrophils, and fibrin in areas of femoral vein thrombi in wild-type mice pretreated with buffer alone, tirofiban, or m-tirofiban. Hr10 was not used since it shows minimal binding to mouse αIIbβ3^16^. In mice pretreated with buffer alone, an immediate and robust accumulation of platelets and fibrin was detected after thrombin application, followed in less than a minute by the increasing accumulation of neutrophils in areas of femoral vein thrombi (**Fig. 4**) (**Supplementary Fig. 7**). Mice pretreated with m-tirofiban reduced the total accumulation of platelets, neutrophils or fibrin as effectively as mice pretreated with tirofiban (**Fig. 4**) (**Supplementary Fig. 7**). There was a significant reduction in neutrophil (but not platelet) accumulation during the first 3 minutes in m-tirofiban-treated animals vs those receiving tirofiban (Supplemental Table 4). The significant reductions in neutrophil and platelet accumulation between the tirofiban- or m-tirofiban-treated group vs. the untreated group remained even during the last 3 minutes of the experiment (Supplemental Table 4).

### Effect of inhibitors on tail vein bleeding and bleeding time

Engagement of platelet αIIbβ3 by cross-linked fibrin drives the dynamic process of clot retraction ^34^, which normally helps consolidate the integrity of the hemostatic plug, restore blood flow, and promote wound closure ^35^. Impaired clot retraction *ex vivo* is reflected by increased bleeding both in mice ^36^ as well as patients receiving any of the three αIIbβ3 partial agonist drugs, abciximab (ReoPro^®^), eptifibatide or tirofiban ^16,37,38^. At the clinical loading dose that suppressed femoral VT, tirofiban caused significant bleeding and prolonged bleeding time. Importantly, mice pretreated with m-tirofiban at the effective anti-VT dose displayed no increase in bleeding and no prolongation of bleeding time (**Fig. 5**).

**Figure 5.**
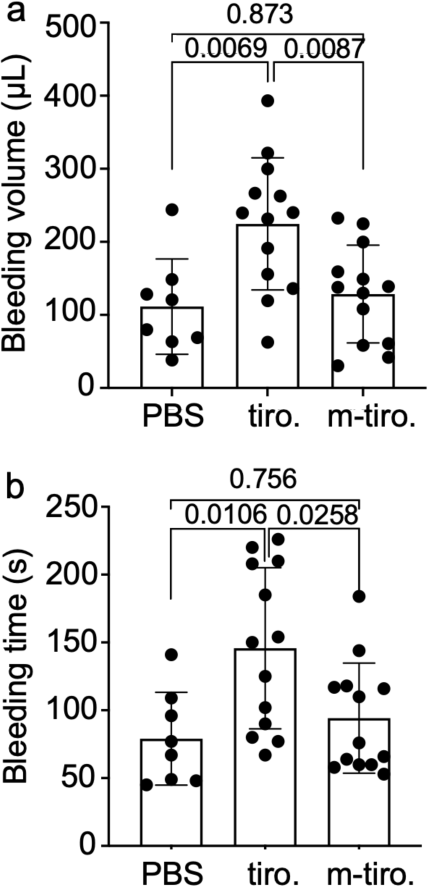
Effect of m-tirofiban and tirofiban on hemostasis in mice. Tail bleeding times were collected for 10 minutes post-tail snip. **a** Bar charts (mean± s.d.) showing baseline bleeding volume in C57BL/6J mice infused intravenously with PBS, tirofiban (tiro.) or m-tirofiban (m-tiro.). **b** Bleeding time in seconds (mean ± s.d.) in the same treated mice. Each dataset was evaluated for normality in GraphPad Prism 10, and subsequently, one-way ANOVA with Tukey’s multiple comparison test was performed.

## Discussion

The data presented in this report establish a direct role for platelet αIIbβ3 in platelet and neutrophil accumulation at site of the activated endothelium of the femoral vein, a primary event in venous thrombogenesis ^7^. These data also demonstrate that a pure αIIbβ3 antagonist (m-tirofiban) effectively suppresses venous thrombogenesis in this thrombosis model without increasing bleeding risk.

That m-tirofiban keeps the native full-length αIIbβ3 in the inactive conformation is revealed in the cryo-EM structure of αIIbβ3 presoaked with 100 μM m-tirofiban. The structure of the complex assumed the canonical bent inactive conformation, a finding supported by the lack of increased expression of the AP5 LIBS epitope by m-tirofiban on ADP- or thrombin-activated human platelet αIIbβ3 (Fig. 3b). A recent study from Morphic Inc., ^39^ reported that a compound, also named m-tirofiban, generated via a different synthetic route, increased expression of the LIBS MBC319.4 epitope on mutationally-activated recombinant αIIbβ3. However, the Morphic Inc. compound has a 20x lower binding affinity for αIIbβ3 and a 33x higher *IC_50 agg_* vs. our m-tirofiban. The poor potency of the Morphic Inc. compound was attributed to high albumin binding. This interpretation is unlikely since our m-tirofiban also exhibits high (>99%) plasma protein binding, a property also shared with 24% of approved drugs ^40–42^, and judged to be of modest relevance to drug efficacy ^43,44^. A possible explanation for the low potency of the Morphic Inc. compound may be structural differences; that is, the ^1^H-NMR spectrum of our m-tirofiban at 400 MHz shows a well-defined aromatic region with clear splitting patterns (**Supplementary Fig. 1d**) in DMSO-d_6_ or CD3OD, whereas the spectrum of the compound reported by Morphic Inc. ^39^ does not display these features.

Pretreatment of wild-type mice with tirofiban at a clinically effective bolus dose significantly reduced the total accumulation of platelets and neutrophils at the site of direct thrombin application to the femoral vein, but the total reduction in fibrin was significant only at a p-value of 0.06 (Fig. 4b). Pretreatment of mice with a bolus dose of m-tirofiban, previously shown effective in preventing photochemical-induced carotid artery thrombosis in humanized mice ^16^, was highly effective in reducing the total accumulation of platelets, neutrophils, and fibrin when compared to the untreated animals (Fig.4b). The total accumulation of each of these parameters was not significantly different from that observed in tirofiban-treated animals (Fig. 4b). The significant reduction in neutrophil (but not platelet) accumulation during the first 3 minutes in m-tirofiban-treated animals vs those receiving tirofiban may reflect a reduced ability of m-tirofiban-bound platelet αIIbβ3 to recruit neutrophils during this period vs. the tirofiban-activated integrin at the tested doses. Whether this observation holds in studies at different drug doses and translates into less thrombosis with m-tirofiban application will require further studies.

At the effective dose that inhibits thrombus initiation, tirofiban-pretreated mice showed increased bleeding and prolonged bleeding times (**Fig. 5**), both of which were not observed in mice pretreated with m-tirofiban. These contrasting outcomes positively correlate with the narrow *IC_50 clot_/IC_50 agg_* ratio of 27.0 for tirofiban and this wide ratio of 1,020 for m-tirofiban. The contribution of albumin binding is likely minor in this system in view of the wide *IC_50 clot_/IC_50 agg_* ratio of 403 for the poor albumin binding of the αIIbβ3 antagonist Hr10, which also preserves hemostasis in the carotid artery thrombosis mouse model ^16^.

A major difference between tirofiban and m-tirofiban is the ability of tirofiban but not m-tirofiban to induce the global activating conformational changes in the native αIIbβ3 receptor (Fig. 2) that have been associated with impaired clot retraction *ex vivo* and increased bleeding in animal models of thrombosis ^16,45^ (this report) and in treated patients with myocardial infarction ^13,46^. A corollary of these observations is that agents that do not induce these activating conformational changes in the native integrin may preserve clot retraction *ex vivo* and hemostasis *in vivo* at the effective anti-thrombosis doses, which is indeed the case with m-tirofiban and Hr10. However, another class of αIIbβ3 antagonists reported by Morphic Inc., termed “closing antagonists,” also prevents the activating conformational changes in αIIbβ3 yet impairs clot retraction *in vitro* ^39^ and hemostasis ^47^. For example, the ligand-mimetic small molecule UR-2922, now discontinued, impairs clot retraction, exhibits a very narrow *IC_50 clot_/IC_50 agg_* ratio of 20.63 (similar to tirofiban), and increases bleeding in pretreated monkeys ^47^. It could be argued that, because of its high affinity to αIIbβ3 (mean *K_d_* of 0.3 nM), UR-2922, like tirofiban (mean *K_d_* of 1.44 nM to active αIIbβ3), is more effective in inhibiting fibrin-mediated clot retraction than m-tirofiban (mean *K_d_* of 23 nM) or Hr10 (mean *K_d_* of 39.7 nM). However, other closing antagonists reported by Morphic Inc. have comparable *K_d_* values to m-tirofiban or Hr10 yet impair clot retraction. For example, the closing inhibitor BMS4-1 (*K_d_* of 54.0 nM to active αIIbβ3) has an *IC_50 clot_* that is ten- and thirty times lower than the respective *IC_50 clot_* for Hr-10 and m-tirofiban ^39^. It thus appears that differences in binding affinity to αIIbβ3 and to albumin are insufficient to explain the differences in the effects of m-tirofiban and Hr10 vs. the “closing antagonist” class on clot retraction.

Of note is that in contrast to m-tirofiban or Hr10, the “closing antagonists” convert not only agonist-activated αIIbβ3 but also the mutationally-activated integrin back to the inactive conformation ^39^, a characteristic of inverse agonists ^48^. Many drugs can act as inverse agonists, but those that failed in clinical trials reversed an intrinsic activity of the respective receptor required to maintain an effective biological function (see, for example ^49^). Previous studies have shown that a sustained occupancy of >95% of the αIIbβ3 binding sites by a drug is required to inhibit clot retraction, implying that a small number of active (fibrin-bound) αIIbβ3 is sufficient for effective clot retraction ^50^. These observations lead us to suggest that for clot retraction, a Goldilocks zone of basal αIIbβ3 activity is required, which is achievable in the presence of m-tirofiban or Hr10, likely related to the unique structural basis of integrin inhibition by these compounds. Vastly shifting the αIIbβ3 conformation in favor of the active (with partial agonists like tirofiban) or the inactive state (with closing antagonists) functionally mimics an absent or non-signaling integrin state, both of which are characterized by impaired clot retraction and bleeding ^51,52^. The present data may thus provide a guiding principle for designing safer inhibitors for other integrins and suggest that m-tirofiban, Hr10, and pure antagonists with a similar structure-activity profile merit further investigations as potential thromboprophylaxis agents in patients at high risk of VT and hemorrhage where anticoagulants are contraindicated.

## Acknowledgments

We thank Drs. K. Alison Rinderspacher and Kelly Dryden for expert technical assistance. We thank Dr. Li Di, Pfizer, Groton, CT, for the helpful discussions. Transmission electron micrographs were recorded at the University of Virginia Molecular Electron Microscopy Core facility (RRID: SCR_019031).

## Author contributions

M.A.A. conceived the project. B.D.A. performed cryo-EM data collection, image processing, and platelet aggregation assays. J-P.X. performed molecular modeling. J.L.A. and J.V.A. performed fibrinogen and AP5 binding studies. B.D.A. and M.A.A. performed clot retraction assays. Synthesis of m-tirofiban was performed by S.X.D. and D.W.L. Plasma protein binding was performed by E.V., J-P.X, and C.B.C. Models of inhibitor binding was performed by D. A. T. Y. G. provided the NCMNP7b polymer and Y.G. and M.Y. provided helpful advice. The venous thrombosis mouse study was carried out by C.F. and M.P., and mouse bleeding by H.S.A. M.P. supervised and analyzed the mouse data. M.A.A. supervised the overall project and wrote the manuscript with input from the co-authors.

## Data and materials availability

All data needed to evaluate the conclusions in the paper are present in the paper and/or the Supplementary Information.

## Code availability

The EM maps and atomic coordinates for the m-tirofiban/full-length integrin complex and the tirofiban-bound integrin will be deposited in the EMDB (www.ebi.ac.uk/pdbe/emdb/) and Protein Data Bank (www.rcsb.org) with accession codes EMDB-1, 2, PDB-1, and EMDB-3, PDB-2 respectively. Full access to these data will be provided upon publication.

## Methods

### Synthesis of (S)-m-tirofiban (m-tirofiban)

#### General procedures

^1^H-NMR and ^13^C-NMR spectra were recorded on an Agilent 400-MR 400-MHz NMR spectrometer. Chemical shifts are reported in parts per million using the residual proton or carbon signal (CD_3_OD: δ_H_ 3.31, δ_c_ 49.00 or CDCl_3_: δ_H_ 7.26, δ_c_ 77.16 (as indicated)) as an internal reference. The apparent multiplicity (s = singlet, d = doublet, t = triplet, q = quartet, m = multiplet, dd = doublet of a doublet, dt = doublet of triplets, qd = quartet of doublets) and coupling constants (in Hz) are reported in that order in the parentheses after the chemical shift. Liquid chromatography and mass spectrometry were performed on a Shimadzu 2020 UFLC mass spectrometer, using a Waters Sunfire column (C18, 5μm, 2.1 mm x 50 mm, a linear gradient from 5 % to 100 % B over 15 min, then 100 % B for 2 min (A = 0.1 % formic acid + H_2_O, B = 0.1 % formic acid + CH_3_CN), flow rate 0.2000 mL/min). All reagents and solvents were used as received from major commercial suppliers, such as Sigma-Aldrich, without further purification. Unless otherwise noted, all air- or moisture-sensitive reactions were run under an argon atmosphere in oven-dried glassware.

### Synthesis of Compound 3 (Supplementary Fig. 1a)

A mixture of Compound 1 (482.9 mg, 1.5 mmol), Compound 2 (563 mg, 1.52 mmol), and cesium carbonate (5.6 g, 17 mmol) in acetone (150 mL) was stirred at 50 ^°^C overnight. The reaction mixture was concentrated under reduced pressure, and the resulting residue was dissolved in ethyl acetate (30 mL). The solution was washed with water (2 x 10 mL), and the solvent was removed *in vacuo*. The residue was purified by flash chromatography on silica gel, eluting with 30 % ethyl acetate in hexane to afford Compound 3 (780 mg, yield 85%) ^53^. LC/MS showed one peak at retention time at 18.0 min with m/z = 611. **^1^H NMR** (400 MHz, CDCl3) δ 7.34 (s, 5H), 7.03 (d, *J* = 8.4 Hz, 2H), 6.78 (d, *J* = 8.4 Hz, 2H), 5.20 (d, *J* = 7.9 Hz, 1H), 5.10 (s, 2H), 4.49 (dd, *J* = 6.1 Hz, *J* = 13.5 Hz, 1H), 4.09 (m, 2H), 3.91 (t, *J* = 6.3 Hz, 2H), 3.01 (d, *J* = 5.5 Hz, 2H), 2.67 (br, 2H), 1.80-1.61 (m, 4H), 1.52 (m, 2H), 1.45 (s, 9H), 1.41 (s, 9H), 1.33-1.24 (m, 3H), 1.15-1.01 (m, 2H). **^13^C-NMR** (101 MHz, CDCl3) δ 170.8, 158.3, 155.8, 155.0, 136.6, 130.6, 129.6, 128.2, 128.0, 114.6, 82.3, 79.3, 67.9, 66.9, 55.5, 44.2, 37.6, 36.4, 36.1, 32.3, 29.6, 28.6, 28.1, 23.3.

### Synthesis of Compound 4 (Supplementary Fig. 1a)

A solution of Compound 3 (780 mg, 1.28 mmol) in 100 mL of methanol in the presence of 10% Pd-C (50 mg) was hydrogenated in a hydrogen atmosphere overnight. The catalyst was removed via filtration over Celite, and the solvent was evaporated *in vacuo* to give a solid (543 mg, yield 90 %). The product was used directly without purification for the next step. LC/MS showed one peak at retention time at 11.0 min with m/z = 477. **^1^H NMR** (400 MHz, CD3OD): δ 7.11 (d, *J* = 8.4 Hz, 2H), 6.82 (d, *J* = 8.4 Hz, 2H), 4.08 (br, 2H), 3.93 (t, *J* = 6.3 Hz, 2H), 3.3 (m, partially overlapped with CD3OD), 2.8 (m, 1H,), 2.66 (br, 2H), 1.75 (m, 2H), 1.71-1.60 (m, 3H), 1.45 (s, 9H), 1.44 (s, 9H), 1.38-1.23 (m, 4H), 1.07 (m, 2H). **^13^C-NMR** (101 MHz, CDCl3): δ 174.6, 158.1, 155.1, 130.5, 129.6, 114.6, 81.3, 79.3, 68.0, 56.6, 44.3, 40.5, 36.4, 36.2, 32.4, 29.7, 28.7, 28.2, 23.4.

### Synthesis of Compound 5 (Supplementary Fig. 1a)

A mixture of Compound 4 (26.7 mg, 0.056 mmol), potassium benzoxazole-2-carboxylate (26 mg, 0.13 mmol), HATU (54 mg, 0.12 mmol), triethylamine (30 mg, 0.3 mmol) in methylene chloride (10 mL) was stirred at room temperature (RT) overnight. The reaction mixture was washed with 1N HCl (2 x 5 mL), saturated sodium bicarbonate (2 x 5 mL), and water (2 x 5 mL). The solvents were removed under reduced pressure. The product was purified by flash chromatography on silica gel, eluting with hexane/EtOAc (4:1) to give Compound 5 as a thick solid (21.4 mg, yield 61.4 %). LC/MS showed one peak at retention time at 14 min with m/z = 622. **^1^H NMR** (400 MHz, CDCl3): δ 7.80 (d, *J* = 7.6 Hz, 1H), 7.72 (d, *J* = 8.0 Hz, 1H), 7.64 (d, *J* = 8.0 Hz, 1H), 7.48 (t, *J* = 7.6 Hz, 1H), 7.42 (t, *J* = 7.6 Hz, 1H), 7.11 (d, *J* = 8.4 Hz, 2H), 6.80 (d, *J* = 8.8 Hz, 2H), 4.93 (dt, *J =* 6.0 Hz, *J* = 13.8 Hz, 1H), 4.05 (br s, 1H), 3.91 (t, *J* = 6.4 Hz, 2H), 3.36 (br s, 1H), 3.18 (qd, *J* = 6.0 Hz, *J* = 14.0 Hz, 2H), 2.66 (br t, *J* = 16.0 Hz, 2H), 1.74 (pentet, *J* = 6.8 Hz, 2H), 1.66-1.61 (m, 3H), 1.48-1.41 (m, 22H), 1.31-1.29 (m, 2H), 1.14-1.03 (m, 2H). **^13^C NMR** (101 MHz, (CDCl3): δ 169.9, 158.4, 155.2, 155.1, 151.3, 140.5, 130.6, 127.6, 125.7, 121.6, 114.7, 112.0, 83.0, 79.3, 68.0, 60.5, 54.3, 37.4, 36.4, 36.1, 29.6, 28.6, 28.2, 23.3, 21.2, 14.3.

### Synthesis and analysis of Compound 6 (m-tirofiban) (Supplementary Fig. 1a-d)

Trifluoroacetic acid (TFA) (1 mL) was added to a solution of Compound 5 (21 mg, 0.033 mmol) in dichloromethane (CH_2_Cl_2_)(5 mL). The mixture was stirred overnight and concentrated *in vacuo*. The resulting residue was triturated with diethyl ether (2 x 3 mL) to give Compound 6 as a white solid (18 mg, TFA salt, yield: 95 %). LC/MS showed one peak at retention time 7.5 min (Fig. S1b) with *m/z* = 466 (Fig. S1c); monitoring at *m/z* 484 confirmed the absence of hydrolysis product(s), observed with exposure to aqueous acid at retention time 7.7 min *m/z* 484 (M+1), 482 (M-1).

**^1^H NMR** (400 MHz, CD3OD): δ 7.84 (dd, *J* = 0.8 Hz, *J* = 8.0 Hz, 1H), 7.73 (d, *J* = 8.0 Hz, 1H), 7.55 (td, *J* = 1.2 Hz, *J* = 7.2 Hz, 1H), 7.48 (td, *J* = 1.2 Hz, *J* = 7.6 Hz, 1H), 7.19 (d, *J* = 8.8 Hz, 2H), 6.81 (d, *J* = 8.4 Hz, 2H), 3.93 (t, *J* = 6.0 Hz, 2H), 3.36 (br s, 1H), 3.34-3.33 (m, 1H), 3.11 (dd, *J* = 7.6 Hz, *J* = 14.8 Hz, 1H), 2.94 (td, *J* = 2.0 Hz, *J* = 12.4 Hz, 2H), 1.93 (br d, *J* = 14.0 Hz, 2H), 1.73 (pentet, *J* = 6.8 Hz, 2H), 1.62-1.55 (m, 1H), 1.54-1.46 (m, 2H), 1.37-1.27 (m, 5H) (Fig. S1d).

**^13^C NMR** (101 MHz, CD3OD): δ 173.9, 159.5, 157.2, 156.5, 152.3, 141.6, 131.3, 130.1, 128.9, 126.9, 122.4, 115.6, 112.7, 68.6, 55.8, 45.3, 37.4, 36.7, 34.8, 30.3, 30.0, 23.9.

### Electron cryomicroscopy (Cryo-EM)

Human full-length αIIbβ3 was purified from outdated platelets as described ^20^. m-tirofiban or tirofiban were added following dialysis at 20-fold molar excess (100 μM m-tirofiban, 72 μM tirofiban) ∼48 hours before EM sample preparation. Cryo-EM data were collected on a Titan Krios electron microscope, and image stacks were motion-corrected during collection using Cryosparc Live as described ^20^.

Processing of m-tirofiban bound αIIbβ3 images was performed in CryoSparc ^54^ on a single collection of 3,133 micrographs (**Supplementary Fig. 2**). Following the CTF assignment, 2,634 micrographs displaying a maximum resolution estimate of 5.2 Å or better were selected for further processing. 2,522,350 particles were selected from templates generated from a previous unliganded αIIbβ3 EM map ^20^, and 737,408 of these were selected for 2D class averaging. 449,891 particles from two consecutive rounds of class averaging and class selection were used for *ab initio* model generation followed by heterogeneous 3D classification against two models. This process was repeated two additional times with three and two models, respectively, retaining the class with the best resolution at each step. The final 3D class containing 112,827 particles subjected to non-uniform refinement yielded a map at 4.1 Å resolution. To determine if any open (hybrid-swingout) form could be identified in the m-tirofiban dataset, we selected particles using templates generated from the open eptifibatide-bound EM map ^20^, which we also employed in particle selection of tirofiban-bound αIIbβ3 particles. 1,086,848 particles were subjected to two rounds of 2D classification followed by five rounds of 3D classification. We were only able to identify a small number of 2D classes containing a total of ∼6,000 particles, which resembled the open form. However, none of these classes had a resolution extending beyond 13 Å.

CryoSparc image processing of tirofiban-bound αIIbβ3 employed 4,229 images (**Supplementary Fig. 4**). During data collection, 975,560 particles were selected in CryoSparc Live from templates generated by a blob picker. Subsequent 2D class averages displayed a number of averages resembling an extended, eptifibatide-like bound species observed in a previous study ^20^, along with a single class resembling the bent conformation seen with unliganded αIIbβ3 in the same study. Subsequently, particles were selected from 3,841 micrographs displaying a maximum resolution estimate of 3.89 Å or better, using templates generated from the eptifibatide-bound EM map ^20^, yielding an initial data set of 2,044,924 particles. 1,448,127 of these were selected from 2D class averages and particles subjected to 8 rounds of 3D classification against two or three models, with a single class retained at each step. The final 3D class contained 97,969 particles and generated a map at 3.9 Å using non-uniform refinement.

### Model building

To build the initial models of the m-tirofiban bound and tirofiban-bound full-length αIIbβ3, the cryo-EM structures of the unliganded full-length αIIbβ3 (8t2v.pdb) and liganded full-length αIIbβ3 (8t2u.pdb) were fit as a rigid body into the respective unsharpened EM maps using PHENIX ^55^. For the m-tirofiban structure, the EM map from refinement with static mask generated from PDB 8t2v was used for TM fitting, EM map from refinement with dynamic mask was used for both ectodomain and ligand structure refinement. m-tirofiban CIF file was generated using PHENIX and used in both refinement and manual adjustment. The models were then improved with iterative real-space refinement in PHENIX and manual adjustment in Coot ^56^. Sharpened maps from PHENIX were also used in manual adjustment. The final models fit the map well with the masked CCs of 0.74 and 0.71, respectively. Structural illustrations were prepared with Chimera or PyMOL.

### Isolation of human platelets

Human blood samples were obtained from healthy volunteers who provided written informed consent under a study that complied with all relevant ethical regulations and was approved by the Human Subjects Committee at the Massachusetts General Hospital in accordance with the Helsinki Principles. All subjects denied taking any medications for at least ten days prior. Blood was drawn directly into 3.2% sodium citrate and used within 3 h. Human platelets were purified from human blood by first centrifugation of the citrated blood samples at 160xg for 10 min at RT to obtain platelet-rich plasma (PRP). Next, PRP was centrifuged at 1,200xg or 1,900xg for 10 min at RT after adding prostaglandin E1 to a final 20 ng/ml concentration. The platelet pellet was washed once in Ringer’s citrate-dextrose buffer (108 mM NaCl, 3.8 mM KCl, 1.7 mM NaHCO3, 21.2 mM sodium citrate, 27.8 mM glucose, 1.1 mM MgCl2, pH 6.5) and resuspended to the same initial PRP volume in supplemented Tyrode’s buffer (127mM NaCl, 5mM KCl, 1.8mM NaH2PO4, 25mM NaHCO3, 10mM dextrose/glucose, pH 7.4 containing 0.02% BSA, 1 mM each of CaCl_2_, and MgCl_2_). To obtain platelet-free plasma (PFP), PRP was centrifuged at 10,000xg for 10 min at RT^57^. Platelets in PRP and after washing for each donor were counted with a hemocytometer.

### Small molecule inhibitor binding to human PFP

Plasma protein binding of m-tirofiban was determined using the rapid equilibrium dialysis (RED) device with tirofiban of known plasma protein binding ^25^, serving as an internal control. 100 μL aliquots of PFP-containing test compound (tirofiban or m-tirofiban at 2 µM final concentration) in triplicate were dialyzed against 300μl PBS in the 50 µl of PFP-containing test/control compounds were removed, mixed with 50μl PBS, and left at room temperature as time-zero samples. The plate was sealed with adhesive film and shaken at 250 rpm on an orbital shaker at 37° C for 24 hours. At the end of the dialysis, aliquots of 50 μL samples from both the buffer and sample side of the dialysis device were removed and mixed with an equal volume of the opposite blank matrix (buffer or plasma) in each sample. All samples were mixed with 400μl of stop solution (acetonitrile containing labetalol at 100ng/ml) and centrifuged at 16,873x g for 5 minutes. Collected samples were submitted for analysis by LC/MS/MS (detailed below). The total recovery of a compound in the experiment was evaluated based on mass balance calculations. The absolute concentration of the initial spiked plasma was not measured but was assumed to be 2 µM based on the amount of compound added and the volume of the solution. Low recovery may indicate nonspecific binding to the RED device or plasma instability issues. Individual measurements were deemed acceptable when the recovery was greater than 75%. The % unbound fraction (*f_u_* (%) was calculated as follows: 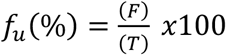, where F is the peak area ratio of analyte/internal standard in the sample on the buffer side of the membrane, and T is the peak area ratio of analyte/internal standard in the sample on the matrix side of the membrane. The area ratios of the test compound to IS in the receiver and donor wells were determined using LC/MS/MS. Recovery was calculated as follows: 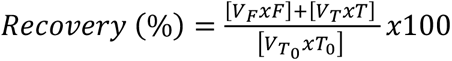, where *V_F_* is the volume of the buffer sample at the beginning of dialysis, *V_T_* is the volume of the matrix sample at the beginning of dialysis, *T_0_* is the compound peak area ratio of analyte/internal standard in the matrix at time zero.

### Liquid Chromatography-Mass Spectrometry/Mass Spectrometry Analysis

Liquid Chromatography-Mass Spectrometry/Mass Spectrometry Analysis. LC-MS/MS analyses were conducted using a Nexera X2 U-HPLC system (Shimadzu Scientific Instruments; Marlborough, MA) coupled to a 7500 QTrap (SCIEX; Framingham, MA). Each analyte was infused to inform the selection of parent/product ion transitions and determine collision energy and cell exit potential values for multiple reaction monitoring (MRM) analyses. Instrument settings were as follows: source temperature, 400°C; curtain gas pressure, 40 psi; ion source gas 1, 45 units; ion source gas 2, 70 units; ion spray voltage, 3 kV; and entrance potential, 10 V. Samples were extracted using 100% acetonitrile containing labetalol as an internal standard. Extracts were injected (10 μL) onto a Zorbax RRHD Eclipse Plus C18 2.1 x 50-mm, 1.8µ column (Agilent, Santa Clara, CA) maintained at 30 °C. The column was eluted at a flow rate of 0.25 mL/min isocratically for 0.5 min at 95% mobile phase A (99.9% water and 0.1% formic acid) followed by a gradient to 95% B (99.9% acetonitrile and 0.1% formic acid) over 2 min, held constant at 95% B for 2 min, and returned to 5% B in 0.2 min. Tirofiban and m-tirofiban eluted between 2.5 and 3.5 min. Positive ion mode MRM transitions were: tirofiban, 441→276 and m-tirofiban, 466→275.

### Binding of inhibitors to ADP-activated platelets

This was quantified by assessing the effects of increasing amounts of tirofiban, m-tirofiban, or Hr10 on displacing Alexa647-conjugated fibrinogen (Alexa647-FB) from ADP-treated platelet αIIbβ3 in 0.02% BSA. 10 µL of increasing concentrations of each compound prepared in Tyrode’s buffer were added to 30µL of washed human platelets in the same buffer and incubated for 20 minutes at RT without stirring. A mixture of Alexa647-FB (at 0.42 or 0.5 µM final concentration) and ADP (at 20 µM) in a total volume of 10µL in Tyrode’s buffer was subsequently added, and the mixture incubated for an additional 30 minutes at RT. The estimated αIIbβ3 in the reaction ranged from 5.42 to 7.99 nM, based on the number of washed platelets for each donor and the estimated 45,000 αIIbβ3 receptors per ADP-activated platelet ^58^. Samples were diluted, fixed in one mL of 2% paraformaldehyde, and analyzed using an LSR Fortessa X-20 flow cytometer (BD Biosciences). Mean fluorescence intensity (MFI) was determined using FlowJo software. Background binding, determined in the presence of 5 mM EDTA, was subtracted. Alexa647-FB binding was expressed as mean fluorescence intensity (MFI) using FlowJo software. Curve-fitting and statistical calculations are performed in SigmaPlot (Systat Software, San Jose, CA). The concentration of the test inhibitor required to induce half-maximal displacement of Alexa647-FB binding (*IC_50_*) was determined and expressed as mean ± s.e.

### LIBS AP5 epitope exposure on αIIbβ3

We compared the effects of m-tirofiban (16 μM), triofiban (0.17μM), and Hr10 (7.6μM), equivalent to 696x, 118x, and 191x the respective *K_d_* to active αIIbβ3) on AP5 epitope expression on ADP-activated washed human platelets. 10µL of Tyrode’s buffer alone or containing tirofiban, m-tirofiban, or Hr10 were added to 30 µL of washed platelets in Tyrode’s buffer containing 1 mM each of Ca^2+^ and Mg^2+^ and 0.02 % BSA and incubated for 20 minutes at RT without stirring. ADP (20µM) and Alexa647-AP5 (10µg/mL) in the same buffer were added in a total volume of 10µL for an additional 30 minutes at RT. Samples were then processed as above for flow cytometry.

### Platelet aggregation

Impedance aggregation measurements were carried out on different days in whole blood from 4-5 young, healthy donors using a Chrono-Log model 700 two-channel lumiaggregation system following the manufacturer’s directions ^16^. Stock dilutions of inhibitor were made in physiologic saline and allowed to solubilize for at least two hours (up to 48 hours) at room temperature. Final dilutions were made in 0.5 mL saline in Chrono-Log cuvettes and warmed at 37°C for 5 min before the addition of 0.5 mL of fresh citrated whole blood. The mixture was warmed for an additional 5 min at 37°C. Measurements were initiated by placing the cuvette in a 37°C chamber with 1,200 rpm stirring with the insertion of a reusable electrode. A one-minute baseline recording is taken, ADP (20µM) is added, and tracing initiated for 6 minutes. Data points for an individual dose curve were serially collected from a single draw and analyzed with SigmaPlot using a least-square fit to a four-parameter logistic curve. The *IC_50 agg_* values were determined from the fitted parameter.

### Clot retraction

The effect of increasing concentrations of inhibitors diluted in saline on clot retraction was measured at RT as described ^16,59^. Digital photographs of the experiment were taken at 15-minute intervals up to 3 hours. Images were analyzed with ImageJ software to determine the areas the clot and plasma occupied. Subsequent data analysis was performed in SigmaPlot. For each inhibitor concentration, a linear regression was fit to the plot of relative clot area against time. The regression slope at each point was plotted against concentration and fitted with a four-parameter logistic curve to determine the inhibitor concentration required to induce half-maximal inhibition of clot retraction (*IC_50 clot_*) (**Supplementary Fig. 6**).

### Models of inhibitor binding to human plasma albumin, platelet αIIbβ3, and fibrinogen

To estimate inhibitor-binding dissociation constants to human albumin (*K_d,A_*) and αIIbβ3 (*K_d,T_*) in the presence of albumin and fibrinogen, sets of Ordinary Differential Equations (ODEs) were constructed to describe each experimental condition. Albumin binding of Hr10, a derivative of hFN10, is considered minimal (1 mM) ^60^. Illustrative models in diagrams are shown in Supplementary Fig.5.

### Model of tirofiban and m-tirofiban binding to platelet-free human plasma (mainly albumin)

A set of ODEs was used to describe the *ex vivo* non-specific binding of tirofiban and m-tirofiban in platelet-free human plasma by equilibrium dialysis (Eqs 1-4).

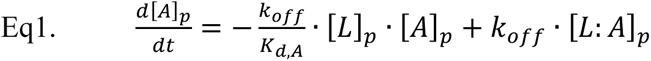

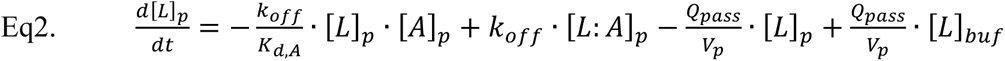

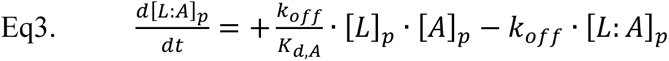

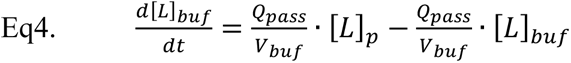

Where [A]*_p_*, [L]*_p_*, [L:A]*_p_*, and [L]*_buf_* are the concentrations of unbound plasma protein (mainly albumin), unbound plasma ligand (tirofiban or m-tirofiban), albumin-bound with ligand, and unbound buffer ligand respectively; *K_d,A_* is the binding dissociation constant of ligand to albumin; *k_off_* is the binding dissociation rate; *Q_pass_* is the passive diffusion rate of ligand across the equilibrium dialysis semipermeable membrane; and *V_p_* and *V_buf_* are the plasma and buffer volumes. The ODEs were simulated to steady state in Berkeley Madonna (v8.3.18), and *K_d,A_* values (448 µM and 6.11µM for tirofiban and m-tirofiban, respectively) fit to achieve experimental unbound fractions (*f_u,p_* of 41.27% and 0.95% for tirofiban and m-tirofiban, respectively, Eq5) in platelet-free plasma in the presence of total albumin and ligand concentrations of 639 µM and 1.0 µM respectively. At steady state, the choice of *k_off_* and *Q_pass_* are irrelevant.

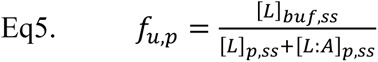

### Model of inhibitor binding to αIIbβ3 in the presence of albumin and fibrinogen

A set of ODEs was used to describe the simultaneous displacement of Alexa647-fibrinogen binding to αIIbβ3 in the presence of tirofiban, m-tirofiban, or Hr10 (Eq 6-12).

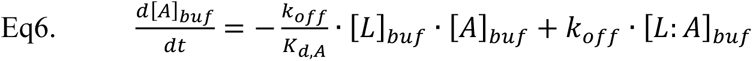

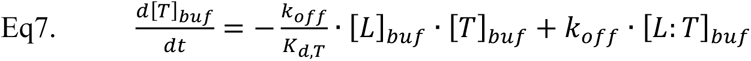

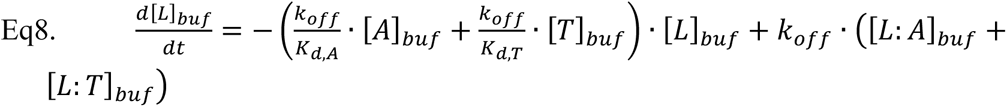

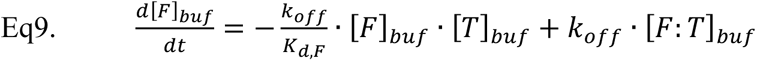

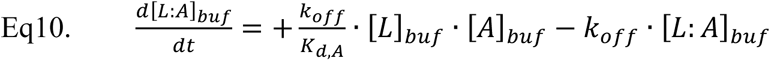

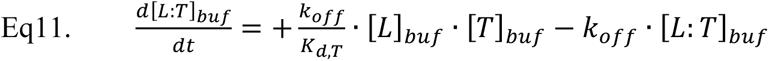

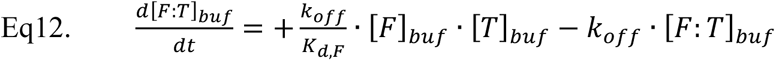

Where [A]*_buf_*, [T]*_buf_*, [L]*_buf_*, [F]*_buf_*, [L:A]*_buf_*, [L:T]_buf_, and [F:T]_buf_ are the concentrations of unbound albumin, unbound αIIbβ3, unbound ligand (tirofiban, m-tirofiban or Hr10), unbound fibrinogen, albumin-bound with ligand, αIIbβ3 bound with ligand, and αIIbβ3 bound with fibrinogen respectively; *K_d,A_* is the binding dissociation constant of ligand to albumin; *K_d,T_* is the binding dissociation constant of ligand to αIIbβ3; *K_d,F_* is the binding dissociation constant of fibrinogen to αIIbβ3; and *k_off_* is the binding dissociation rate.

The ODEs were simulated to steady state in Berkeley Madonna (v8.3.18), and mean *K_d,T_* values of 1.44 nM, 23 nM, and 39.7 nM for tirofiban, m-tirofiban, and Hr10, respectively, fit to achieve experimental relative Alexa647-fibrinogen displacement by respective inhibitor mean *IC_50_* values of 9.95 nM, 138 nM, and 175 nM (*DIS_F:T_*, Eq15) in the presence of 3 µM total albumin (0.02% BSA), 420 nM or 500 nM total Alexa647-fibrinogen, and individual experimental total αIIbβ3 (∼8 nM). Relative fibrinogen displacement by ligand is a function of the occupancy of fibrinogen to αIIbβ3 in the presence (*OCC_F:T_*, Eq13) and absence (*OCC’_F:T_*, Eq14) of ligand. The *K_d,A_* of ligands to the incubation 0.02% BSA was assumed equivalent to that estimated to human albumin. The *K_d,F_* of fibrinogen to αIIbβ3 was assumed to be 143 nM ^58^. At steady state, the choice of *k_off_* is irrelevant, and fibrinogen occupies 74% of αIIbβ3 in the absence of ligand.

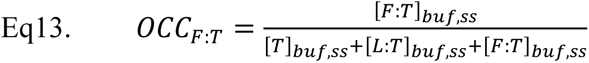

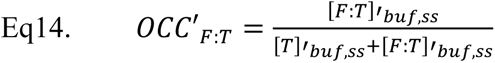

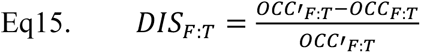

### Model of inhibitor binding to αIIbβ3 and albumin

A set of ODEs was used to describe the simultaneous binding of tirofiban, m-tirofiban, and Hr10 to αIIbβ3 and albumin in 1:1 diluted whole blood prior to the addition of ADP (Eq 16-21).

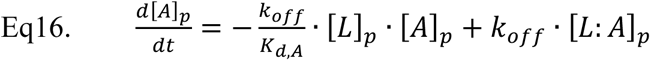

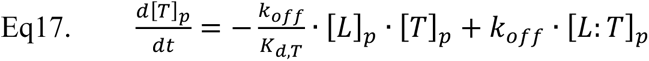

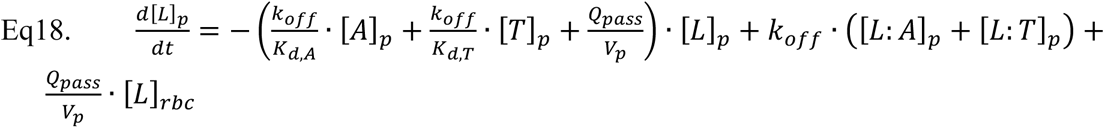

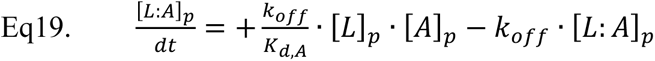

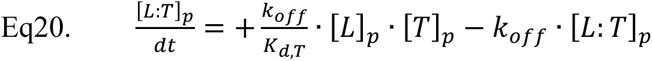

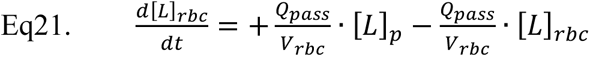

Where [A]*_p_*, [T]*_p_*, [L]*_p_*, [L:A]*_p_*, [L:T]_p_, and [L]*_rbc_* are the concentrations of unbound plasma albumin, unbound plasma αIIbβ3, unbound plasma ligand (tirofiban, m-tirofiban or Hr10), albumin-bound with ligand, αIIbβ3 bound with ligand, and unbound ligand in red blood cells respectively; *K_d,A_* is the binding dissociation constant of ligand to albumin; *K_d,T_* is the binding dissociation constant of ligand to αIIbβ3; *k_off_* is the binding dissociation rate; *Q_pass_* is the passive diffusion rate of ligand in and out of red blood cells; and *V_p_* and *V_rbc_* are the plasma and red blood cell volumes. *K_d,T_* of tirofiban to inactive αIIbβ3 are 15 nM ^28^ and 1.44 nM (this report). *K_d,T_* of m-tirofiban and Hr10 to inactive or active αIIbβ3 are 23 nM and 39.7 nM, respectively. The ODEs were simulated to a steady state in Berkeley Madonna (v8.3.18) to estimate ligand occupancy of αIIbβ3 (*OCC_L:T_*, Eq22). At steady state, the choice of *k_off_* and *Q_pass_* are irrelevant. Non-specific binding of ligand to red blood cells was assumed to be negligible. In this assay, the 1:1 dilution of whole blood with buffer dilutes plasma albumin to 227 µM and αIIbβ3 to 14.5 nM (average 45K receptors/platelet and average 300K platelets per µL undiluted blood), assuming a 45% hematocrit.

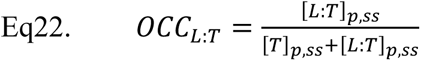

### Femoral vein thrombosis model

Intravital confocal imaging using multiple fluorescent probes was employed to image thrombus formation in real-time in living mice as described ^31^. In this model, direct application of thrombin to the exposed vein precleaned of connective tissue allows rapid endothelial activation with von Willebrand factor and P-selectin surface exposure to drive thrombus initiation, as there was no clot formation when these two proteins are lacking ^31^. Briefly, C57BL/6J mice (6-16 weeks old) of similar weight (untreated: 25.4 ± 1.2g; tirofiban: 24.9 ± 1.1g; m-tirofiban: 25.3 ± 0.8g, mean ± 1 sd), 60% males and 40% females, were assigned to groups regardless of sex or age. All groups had the same criteria for mouse selection. The numbers in each group were not tracked for these demographics. The mouse numbers were predetermined before the studies, were not guided by data accumulation, and were based on prior studies that showed significant differences using these same *in vivo* systems in our laboratory. All experiments were done blinded by the operator as to the outcome until the data was subsequently analyzed. Mice were anesthetized with an intraperitoneal injection of Nembutal (80 mg/kg) beginning 30 minutes before surgery. Mice were also maintained on a 37°C heating pad. Anti-Ly6G F(ab′)_2_ (0.3 µg/g mouse, BD Bioscience), the nonfunctional anti-CD41 F(ab′)_2_ (0.3 µg/g mouse, clone MWReg30, BD Bioscience ^61^, and anti-fibrin (0.3 µg/g mouse ^62^ were infused as 100-μL boluses via a catheter placed into the jugular vein to detect neutrophils, platelets, and fibrin, respectively. Antibodies were labeled using Alexa-fluor mAb labeling kits from Invitrogen (Carlsbad, CA). A skin flap extending from the left ankle to the thigh was removed to expose the intact femoral vein; the vessel was cleared of connective tissue, avoiding damage, and was clearly visualized under a 10X objective with a clear gel (GenTeal Tears Lubricant Eye Gel, Alcon) between the objective lens and the vessel. Tirofiban and m-tirofiban were each diluted in 100 μl of PBS, kept at RT for ∼5 minutes, and then infused through the jugular vein cannula into mice over 5 minutes. Tirofiban bolus dose was 0.025μg/g, equivalent to the clinical loading dose of 25 μg/kg (https://www.drugs.com/pro/tirofiban-injection.html#s-34068-7), effectively blocked photochemical-or ferric chloride-induced carotid artery thrombosis in humanized and WT mice, respectively ^16,45^. m-tirofiban bolus dose was 0.3 μg/g, previously shown to effectively inhibit photochemical-induced injury of the carotid artery in humanized mice ^16^. Confocal time-lapse videos of neutrophils rolling along the venous endothelium were captured pre-(baseline) and 5 minutes post-drug infusion. The exposed intact femoral vein was dabbed and dried with Kim wipes, and one µL solution of human α-thrombin (9 mg/ml) (Hematologic Technologies, Inc.) was applied directly onto the surface of the femoral vein for 2 seconds. The clear gel was quickly reapplied, and time-lapse intravital fluorescence microscopy was initiated immediately to monitor thrombus formation after thrombin application. Animals were then euthanized and disposed of. Confocal time-lapsed images of platelet, neutrophil, and fibrin accumulation were collected at 50 frames/min for 15 minutes. Fluorescence per µm2 was determined from the videos and analyzed using Slidebook 6.0 software (Intelligent Imaging Innovations) to calculate the area occupied by platelets, neutrophils, and fibrin for each study. The area under the curve (AUC) for each animal was calculated, the mean and se for each of the three groups quantified, and data analyzed using One-way ANOVA with Tukey’s multiple comparisons test in GraphPad Prism 10. All mouse experiments complied with all relevant ethical regulations and were approved by the IACUC of the CHOP, and all investigators adhered to NIH guidelines for the care and use of laboratory animals.

### Tail vein bleeding model

This was performed as described ^16^. 8-week-old C57BL/6J mice of similar weight, age, and sex in each were anesthetized with sodium pentobarbital (80 mg/kg) 30 minutes prior to surgery. All experiments were done blinded by the operator as to the outcome until the data was subsequently analyzed. Mice received intravenous PBS, tirofiban, or m-tirofiban (in the dilutions and amounts used in the vein thrombosis model). After 5 min, the tip of the mouse tail (8 mm) was amputated with a sharp razor blade, and total blood loss and total bleeding time were quantified. Blood loss was limited to only 10 minutes post-tail snip. Over this timeframe, insufficient blood loss occurred to consider limiting blood loss. All mice were sacrificed at the end of the bleeding study.

### Statistical calculations

The data points from each replicate from fibrinogen displacement and whole blood aggregation experiments were individually fit to a sigmoidal function to determine the *IC_50_*, Hill coefficient, and the minimum and maximum values. Individual *IC_50_* values for fibrinogen displacement were used to calculate *K_d_* values for each replicate using ODE equations 6-15, reported as the geometric mean of the replicates. Data from the aggregation experiments were scaled to a maximum of 1.0 and a minimum of 0 using the minimum and maximum values from the individual fits and were combined and again fit to a four-parameter sigmoidal curve using non-linear regression to determine parameters and the respective standard errors. Data points for clot retraction were combined without scaling. To evaluate variability between subjects in each group, logarithms of *IC_50_*s (in nM) are derived for each subject, and the mean and associated parameters are determined for each group and shown in the Supplementary Table. 3. Comparisons of the AUC values from the thrombogenesis experiments and bleeding analyses are analyzed without transformation. Analyses are performed with GraphPad Prism 10. Differences were considered statistically significant when p-values were < 0.05.

## Supplementary Information

**Supplementary Figure 1.**
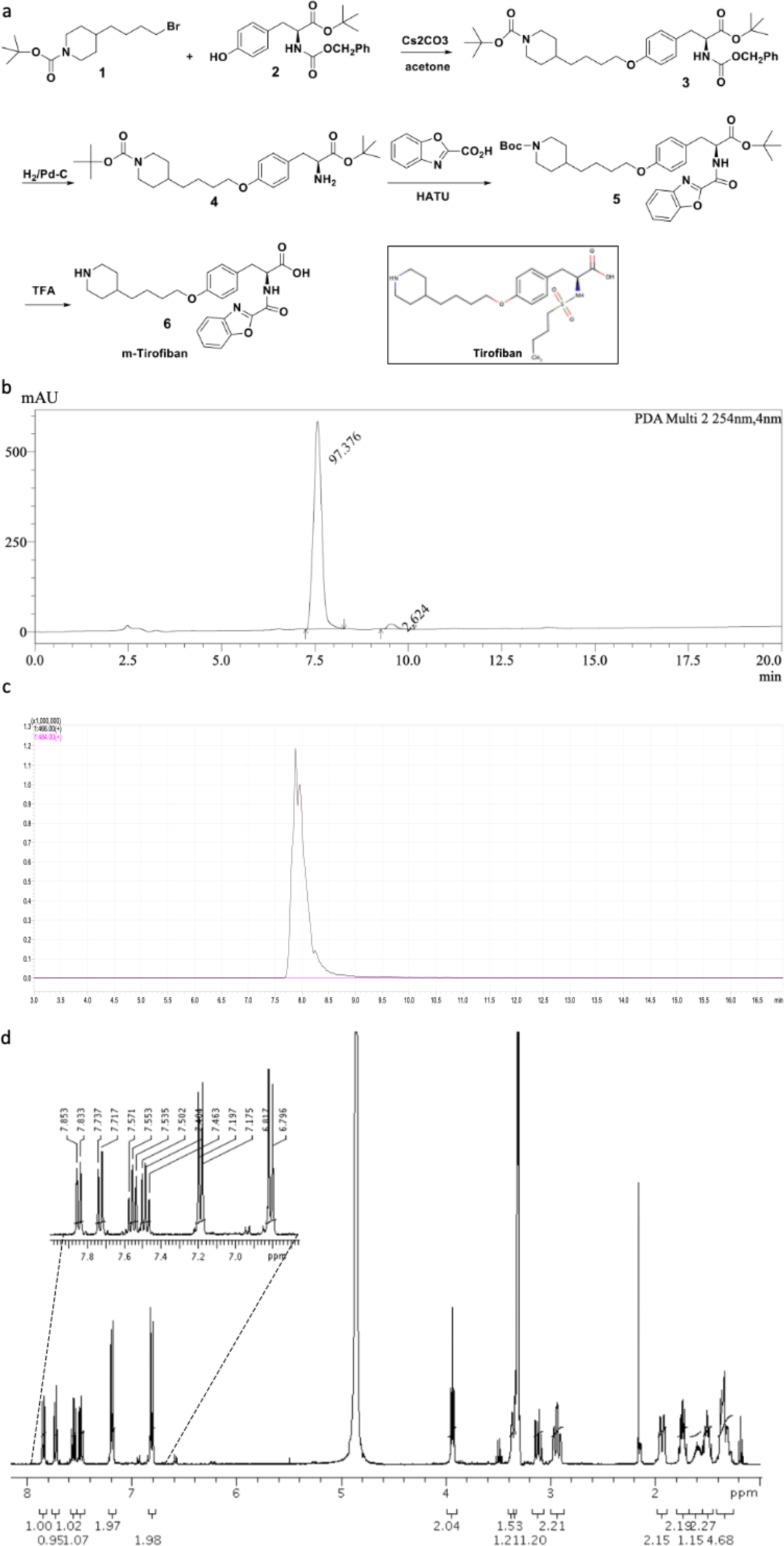
Synthesis and analysis of m-tirofiban. **a** Synthetic scheme. Tirofiban’s chemical structure is shown for comparison. **b** PDA chromatogram of m-tirofiban (y-axis: intensity (mAU); x-axis: time (min); **c** Mass chromatogram of m-tirofiban (y-axis: intensity (x 1,000,000); x-axis: time (min); **d** ^1^H NMR spectrum of m-tirofiban (CD_3_OD: δ_H_ 3.31 ppm; H_2_O: δ_H_ 4.87 ppm) (x-axis = δ scale (ppm)); inset: aromatic region of m-tirofiban.

**Supplementary Figure 2.**
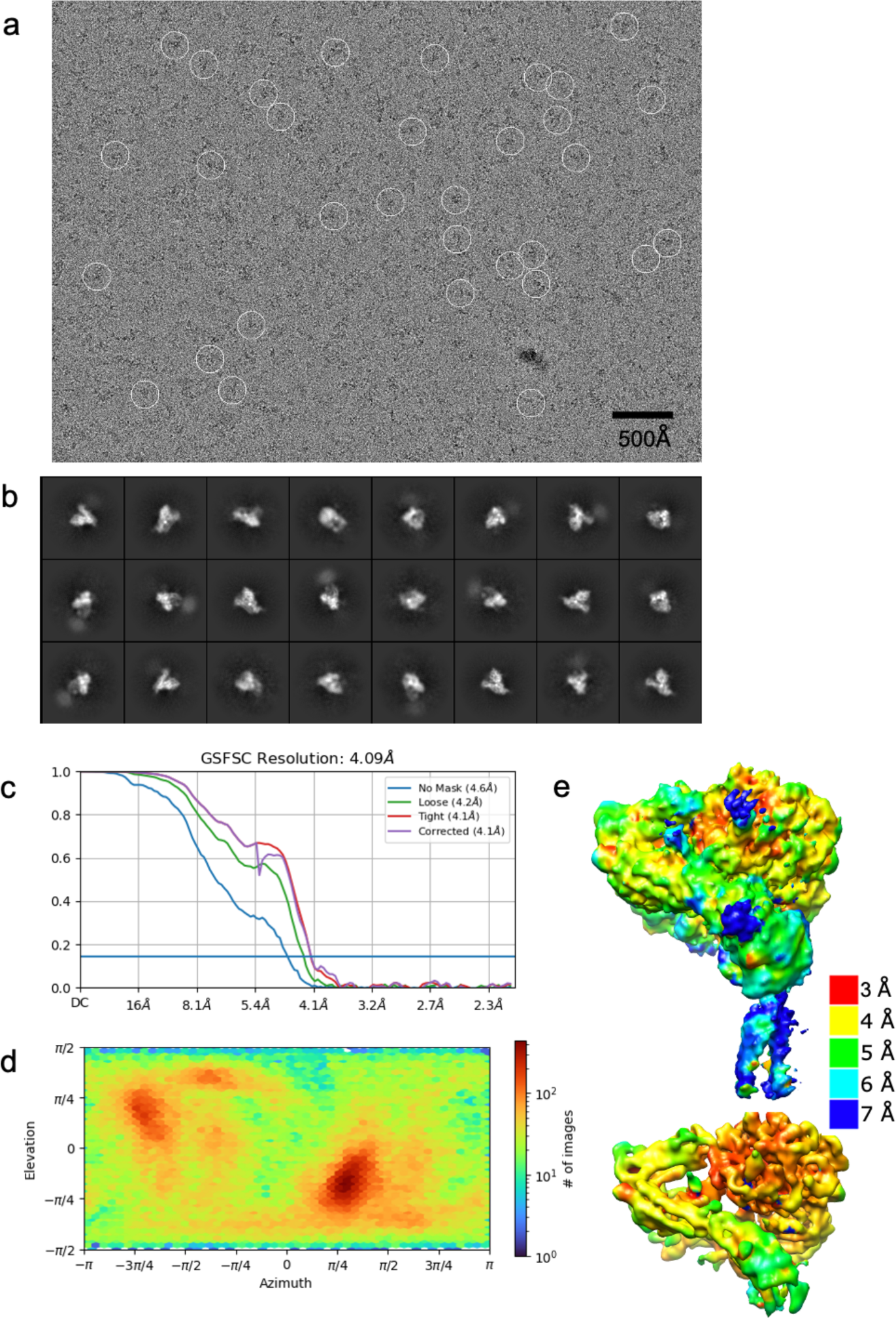
EM processing of full-length αIIbβ3/m-tirofiban complex. **a** A representative cryo-EM micrograph. Selected particles used in the final structure are indicated with white circles. The scale bar at the lower right shows 500 Å. **b** Reference-free class averages generated from selected particles. The averages clearly show diffuse density for the transmembrane regions. **c** Fourier-shell correlation curves for the two half maps for the final round of refinement. The 0.143 cutoff criterion has been used to determine the resolution. **d** Heat map showing the orientation distribution of particles. Euler angles are indicated along the X and Y coordinates. Color encoding for particle numbers is shown at the right. **e** Color-coded local resolution map generated using the 0.143 cutoff criterion. The top and bottom figures show different isosurface values. Colors encode graded resolutions between 3.0 Å (red) and 7.0 Å (blue).

**Supplementary Figure 3.**
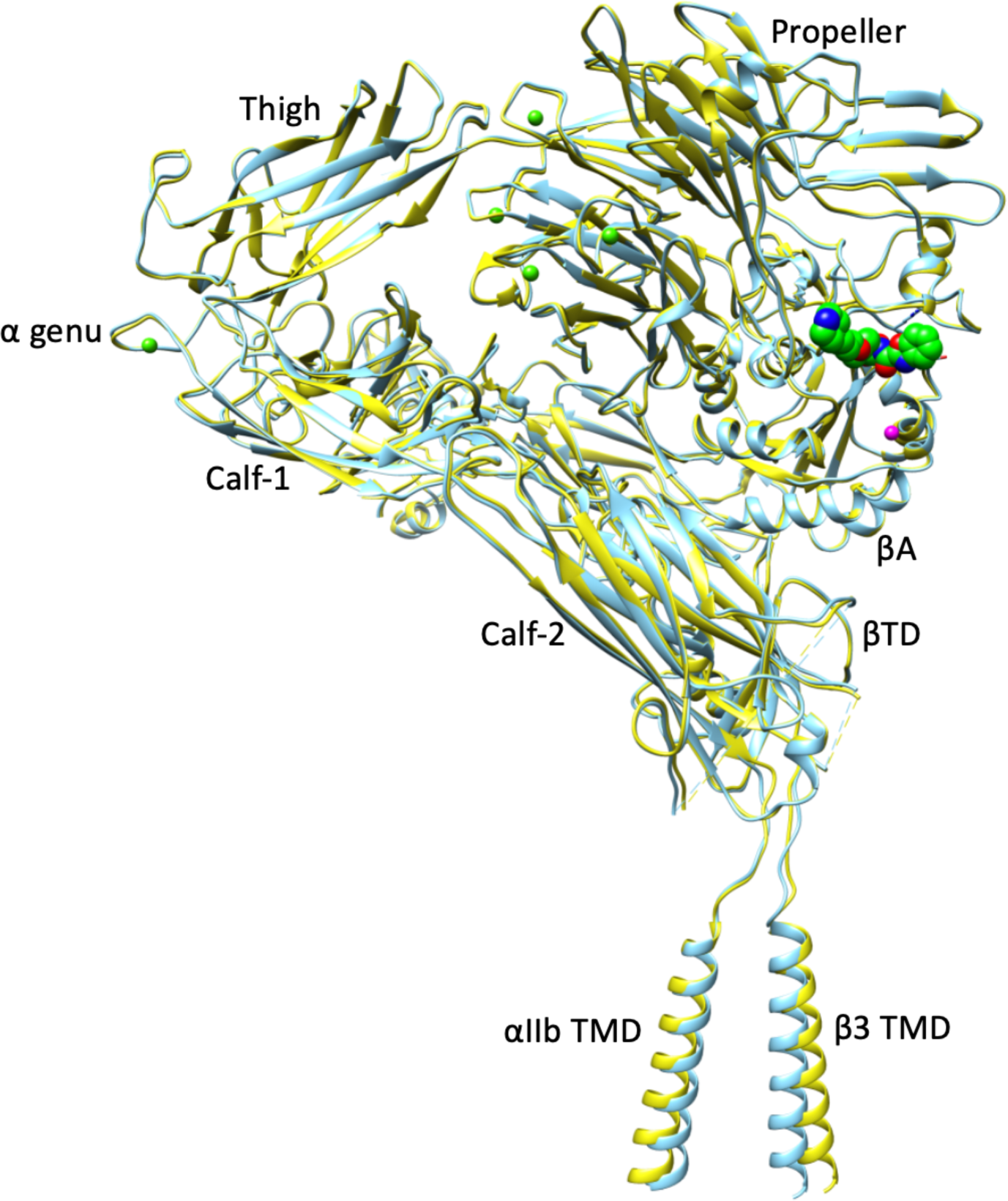
Cryo-EM structures of unliganded and m-tirofiban-bound full-length αIIbβ3. Ribbon diagrams of the superposed structures of unliganded (in yellow)- and m-tirofiban-bound (light blue) full-length αIIbβ3 shown in the same orientation as in Fig. 1a. m-tirofiban atoms are shown as spheres with carbons in green, oxygens in red, and nitrogens in blue. The Ca^2+^ at ADMIDAS is shown as a magenta sphere. The four metal ions (in green spheres) at the bottom of the propeller and the one at the α-genu are shown as green spheres.

**Supplementary Figure 4.**
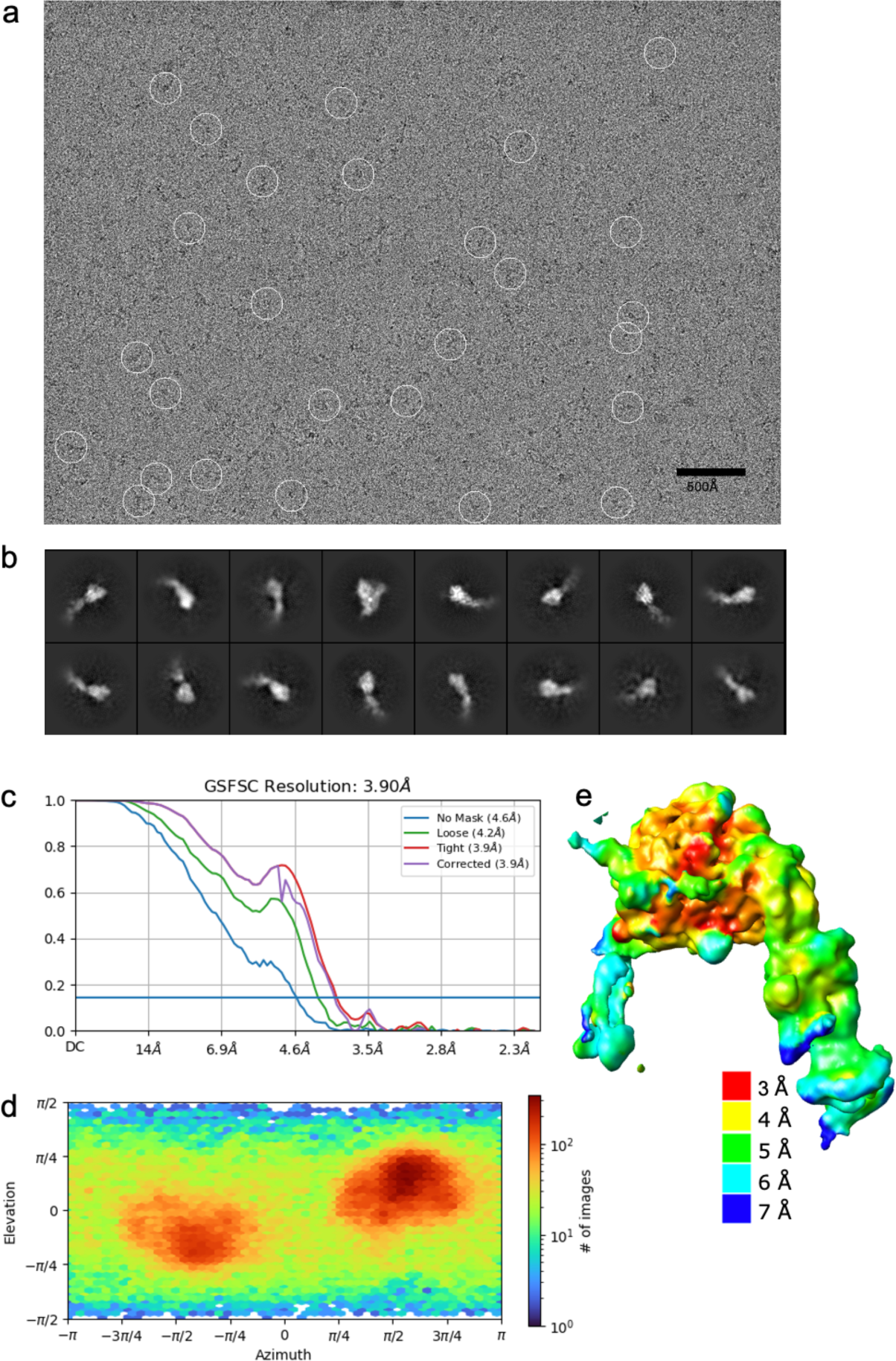
EM processing of full-length αIIbβ3/tirofiban complex. **a** A representative cryo-EM micrograph. Selected particles used in the final structure are indicated with white circles. The scale bar at the lower right shows 500Å. **b** Reference-free class averages generated from selected particles. The majority are not in the bent conformation, but a small number are (example shown in the top row, fourth from left). Domains attributable to regions beyond the integrin headpiece in the extended conformation cannot be identified in the averages. **c** Fourier-shell correlation curves for the two half maps for the final round of refinement. The 0.143 cutoff criterion has been used to determine the resolution. **d** Heat map showing the orientation distribution of particles. Euler angles are indicated along the X and Y coordinates. Color encoding for particle numbers is shown at the right. The distinct red patches indicate a strong preferential orientation for two orientations separated by 180° in azimuth. **e** A color-coded local resolution map of the integrin is generated using the 0.143 cutoff criterion. Colors encode graded resolutions between 3.0 Å (red) and 7.0 Å (blue).

**Supplementary Figure 5.**
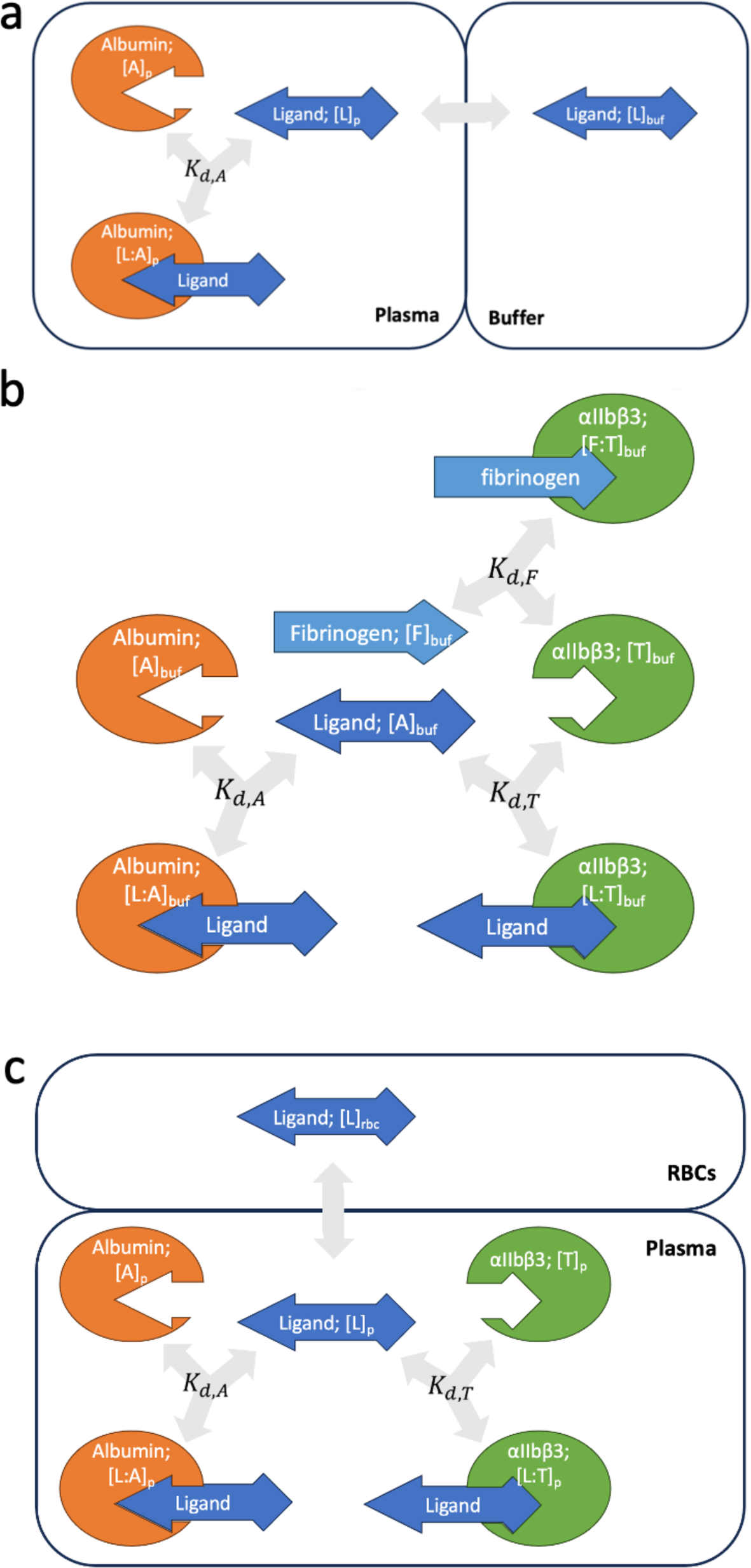
Diagrams of the ODE models used. **a** Diagram of tirofiban and m-tirofiban binding to platelet-free human plasma (Eq 1-5). **b** Diagram of inhibitor binding to αIIbβ3 in the presence of albumin and fibrinogen (Eq 6-15). **c** Diagram of inhibitor binding to αIIbβ3 and albumin (Eq 16-22).

**Supplementary Figure 6.**
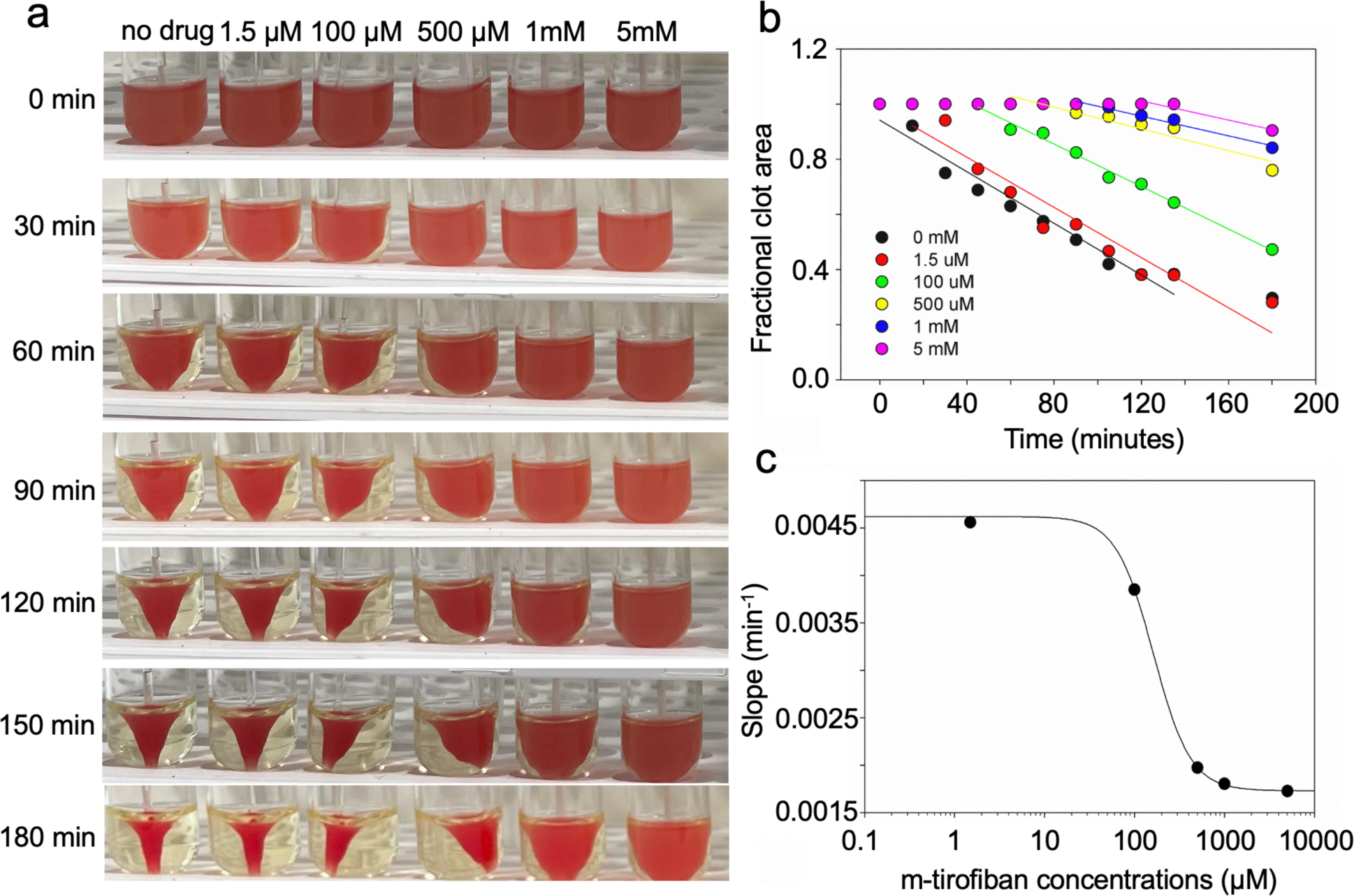
Representative clot retraction titration and analysis. **a** Kinetics of clot retraction in the presence of varying concentrations of m-tirofiban in one experiment. Clot retraction took place around a central glass rod. A total of 5 μl of red blood cells were added per 1 ml reaction to enhance color contrast for photography. Photographs shown were taken at timed intervals (in minutes) after the addition of thrombin. **b** Time course of the titration shown in panel (a). The plot shows the fractional area occupied by the clot as a function of time, with a linear regression through the points. Points from the lag period noted at higher m-tirofiban concentrations were not included in the linear regression. **c** Plot (black circles) of the absolute value of each slope in panel (b) as a function of the m-tirofiban concentration. The points have been fit to a four-parameter, sigmoidal function with non-linear regression.

**Supplementary Figure 7.**
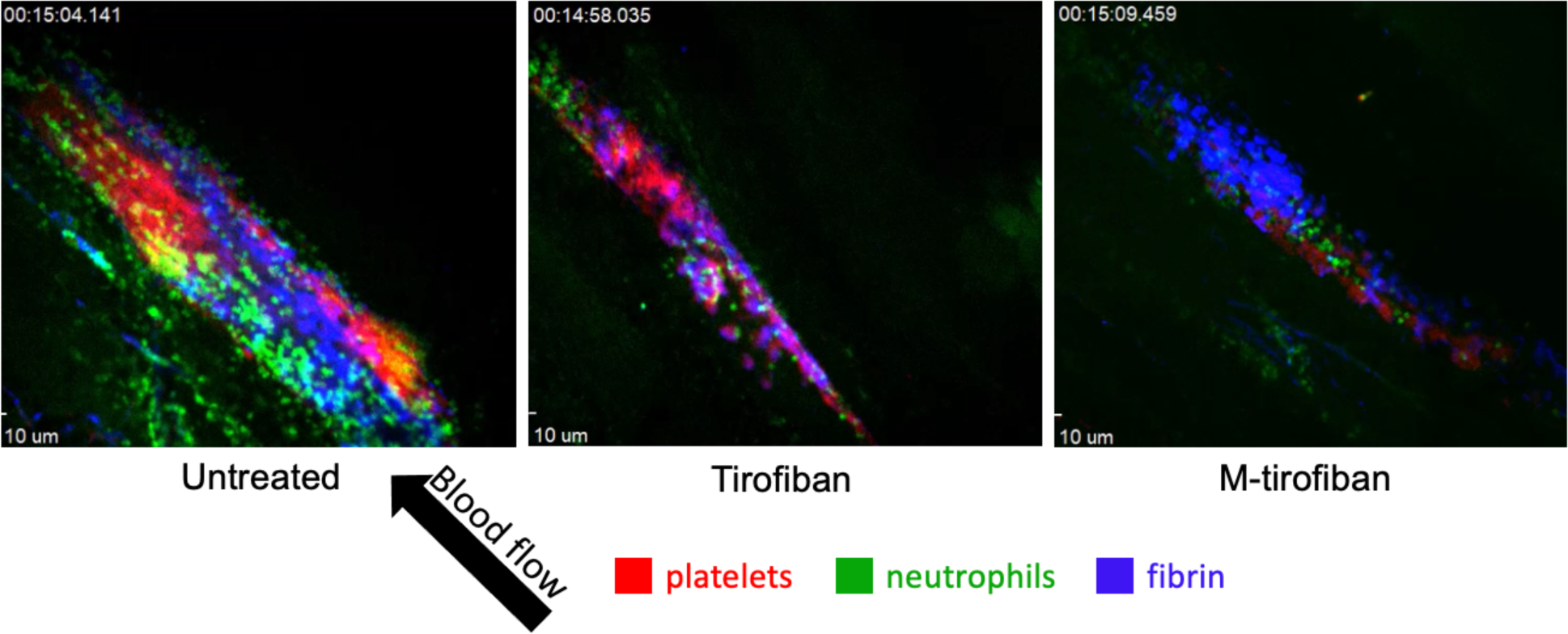
Images of femoral vein thrombogenesis. Representative images at ∼15 minutes after a thrombin activation of the endothelium was established in a femoral vein (see recorded time in each image) in untreated (left), tirofiban-treated (middle), and m-tirofiban treated (right) vessels. Size bars (Scale 10μm) are included in the bottom left of each figure, and the direction of blood flow is indicated by the large black arrow. Incorporated platelets are red, while adherent/rolling neutrophils are green, and fibrin is blue.

**Supplementary Table 1.** Cryo-EM data collection, refinement, and validation statistics.

|  | (EMDB-1); (EMDB-2)<br>(PDB-1) | (EMDB-3)<br>(PDB-2) |
| --- | --- | --- |
| *Data collection and processing* |  |  |
| Magnification | 81,000 | 81,000 |
| Voltage (kV) | 300 | 300 |
| Electron exposure (e-/Å<sup>2</sup>) | 50 | 50 |
| Defocus range (μm) | -2.2 to -1.2 | -2.2 to -1.2 |
| Pixel size (Å) | 1.08 | 1.08 |
| Symmetry imposed | C1 | C1 |
| Initial particle images (no.) | 2,522,350 | 4,245,069 |
| Final particle images (no.) | 112,827 | 97,969 |
| Map resolution (Å) | 4.1 | 3.9 |
| FSC threshold | 0.143 | 0.143 |
| Map resolution range (Å) | 2.9 – 8.6 | 2.8 – 7.5 |
| *Refinement* |  |  |
| Initial model used (PDB code) | 8t2v | 8t2u |
| Model composition |  |  |
| Non-hydrogen atoms | 13,148 | 7,364 |
| Protein residues | 1,664 | 925 |
| Ligands | 1 | 1 |
| *B* factors (Å<sup>2</sup>) |  |  |
| Protein | 258 | 280 |
| Ligand | 285 | 241 |
| R.m.s. deviations |  |  |
| Bond lengths (Å) | 0.002 | 0.002 |
| Bond angles (°) | 0.409 | 0.425 |
| Validation |  |  |
| MolProbity score | 1.39 | 1.57 |
| Clashscore | 3.94 | 5.89 |
| Poor rotamers (%) | 0.99 | 0.9 |
| Ramachandran plot |  |  |
| Favored (%) | 96.68 | 96.3 |
| Allowed (%) | 3.26 | 3.7 |
| Disallowed (%) | 0.06 | 0 |

**Supplementary Table 2.** Human plasma protein binding of tirofiban and m-tirofiban.

| Sample Name | Batch No. | Human platelet-free plasma |  |  |  |  |
| --- | --- | --- | --- | --- | --- | --- |
|  |  | % Unbound (n = 3) | SD | % Bound | % Recovery (n = 3) | SD |
| tirofiban | N/A | 41.27 | 3.03 | 58.73 | 116.14 | 4.51 |
| m-tirofiban | N/A | 0.95 | 0.21 | 99.05 | 94.45 | 4.07 |

**Supplementary Table 3.** Mean, s.d., and s.e. of logged *IC_50_* values for the donors presented in Figures 3a, c, and d.

| Experiment | FB displacement |  |  | Platelet aggregation |  |  | Clot retraction |  |  |
| --- | --- | --- | --- | --- | --- | --- | --- | --- | --- |
| Treatment Group | tiro. | m-tiro. | Hr10 | tiro. | m-tiro. | Hr10 | tiro. | m-tiro. | Hr10 |
| mean | 1.0232 | 2.1392 | 2.2429 | 0.7826 | 1.9659 | 1.7654 | 2.2401 | 5.0147 | 4.3886 |
| s.d. | 0.1060 | 0.1555 | 0.1050 | 0.1455 | 0.1363 | 0.1166 | 0.3217 | 0.0735 | 0.1190 |
| s.e. | 0.0612 | 0.0898 | 0.0606 | 0.0728 | 0.0609 | 0.0583 | 0.1857 | 0.0425 | 0.0687 |
| 95% confidence limit | 0.2627 | 0.3856 | 0.2604 | 0.2315 | 0.1692 | 0.1855 | 0.7974 | 0.1823 | 0.2950 |

**Supplementary Table 4.** Early and late accumulation of cells and fibrin in thrombi.

| **Platelet AUC (First 3 minutes)** |  |  |  |
| --- | --- | --- | --- |
| One-way ANOVA* | Mean of difference | 95% CI of difference | Individual P-value |
| Untreated vs. tiro. | 78393 | 19115 to 137670 | 0.0092 |
| Untreated vs. m-tiro. | 85969 | 31366 to 140572 | 0.0023 |
| tiro. vs. m-tiro. | 7576 | -44018 to 50171 | 0.9251 |
| **Neutrophil AUC (First 3 minutes)** |  |  |  |
| Untreated vs. tiro. | 3051 | -12401 to 18502 | 0.8692 |
| Untreated vs. m-tiro. | 19143 | 4910 to 33375 | 0.0081 |
| tiro. vs. m-tiro. | 16092 | 2644 to 29541 | 0.0181 |
| **Fibrin AUC (First 3 minutes)** |  |  |  |
| Untreated vs. tiro. | 48345 | -5916 to 102605 | 0.0853 |
| Untreated vs. m-tiro. | 37741 | -12240 to 87722 | 0.1588 |
| tiro. vs. m-tiro. | -10604 | -57832 to 36624 | 0.8346 |
| **Platelet AUC (Last 3 minutes)** |  |  |  |
| Untreated vs. tiro. | 61115 | 7353 to 114876 | 0.0248 |
| Untreated vs. m-tiro. | 72423 | 22901 to 121944 | 0.0043 |
| tiro. vs. m-tiro. | 11308 | -35485 to 58102 | 0.8113 |
| **Neutrophil AUC (Last 3 minutes)** |  |  |  |
| Untreated vs. tiro. | 31273 | 9790 to 52756 | 0.0045 |
| Untreated vs. m-tiro. | 34649 | 14861 to 54438 | 0.0009 |
| tiro. vs. m-tiro. | 3377 | -15322 to 22075 | 0.8892 |
| **Fibrin AUC (Last 3 minutes)** |  |  |  |
| Untreated vs. tiro. | 46368 | 4808 to 87927 | 0.0277 |
| Untreated vs. m-tiro. | 43484 | 5202 to 81766 | 0.0249 |
| tiro. vs. m-tiro. | -2884 | -39057 to 33289 | 0.9772 |

\* Data analysis used one-way ANOVA with Tukey's multiple comparison test. CI, confidence interval.

## References

1 Heit, J. A. Epidemiology of venous thromboembolism. Nat Rev Cardiol 12, 464–474, doi:10.1038/nrcardio.2015.83 (2015).

2 Schulman, S., Hu, Y. & Konstantinides, S. Venous Thromboembolism in COVID-19. Thromb Haemost 120, 1642–1653, doi:10.1055/s-0040-1718532 (2020).

3 Khan, F., Tritschler, T., Kahn, S. R. & Rodger, M. A. Venous thromboembolism. Lancet 398, 64–77, doi:10.1016/S0140-6736(20)32658-1 (2021).

4 Prandoni, P. et al. An association between atherosclerosis and venous thrombosis. N Engl J Med 348, 1435–1441, doi:10.1056/NEJMoa022157 (2003).

5 Wolberg, A. S. et al. Venous thrombosis. Nat Rev Dis Primers 1, 15006, doi:10.1038/nrdp.2015.6 (2015).

6 Sevitt, S. The structure and growth of valve-pocket thrombi in femoral veins. J Clin Pathol 27, 517–528, doi:10.1136/jcp.27.7.517 (1974).

7 von Bruhl, M. L. et al. Monocytes, neutrophils, and platelets cooperate to initiate and propagate venous thrombosis in mice in vivo. J Exp Med 209, 819–835, doi:10.1084/jem.20112322 (2012).

8 Becattini, C. et al. Aspirin for preventing the recurrence of venous thromboembolism. N Engl J Med 366, 1959–1967, doi:10.1056/NEJMoa1114238 (2012).

9 Brighton, T. A. et al. Low-dose aspirin for preventing recurrent venous thromboembolism. N Engl J Med 367, 1979–1987, doi:10.1056/NEJMoa1210384 (2012).

10 Simes, J. et al. Aspirin for the prevention of recurrent venous thromboembolism: the INSPIRE collaboration. Circulation 130, 1062–1071, doi:10.1161/CIRCULATIONAHA.114.008828 (2014).

11 Cavallari, I. et al. Frequency, Predictors, and Impact of Combined Antiplatelet Therapy on Venous Thromboembolism in Patients With Symptomatic Atherosclerosis. Circulation 137, 684–692, doi:10.1161/CIRCULATIONAHA.117.031062 (2018).

12 Nurden, A. T. & Nurden, P. GPIIb/IIIa antagonists and other anti-integrins. Semin Vasc Med 3, 123–130, doi:10.1055/s-2003-40670 (2003).

13 Ley, K., Rivera-Nieves, J., Sandborn, W. J. & Shattil, S. Integrin-based therapeutics: biological basis, clinical use and new drugs. Nat Rev Drug Discov 15, 173–183, doi:10.1038/nrd.2015.10 (2016).

14 Honda, S. et al. Topography of ligand-induced binding sites, including a novel cation-sensitive epitope (AP5) at the amino terminus, of the human integrin beta 3 subunit. J Biol Chem 270, 11947–11954 (1995).

15 Van Agthoven, J. F. et al. Structural basis for pure antagonism of integrin alphaVbeta3 by a high-affinity form of fibronectin. Nat Struct Mol Biol 21, 383–388, doi:10.1038/nsmb.2797 (2014).

16 Adair, B. D. et al. Structure-guided design of pure orthosteric inhibitors of alphaIIbbeta3 that prevent thrombosis but preserve hemostasis. Nat Commun 11, 398, doi:10.1038/s41467-019-13928-2 (2020).

17 Xu, X. P. et al. Three-Dimensional Structures of Full-Length, Membrane-Embedded Human alpha(IIb)beta(3) Integrin Complexes. Biophys J 110, 798-809, doi:10.1016/j.bpj.2016.01.016 (2016).

18 Choi, W. S., Rice, W. J., Stokes, D. L. & Coller, B. S. Three-dimensional reconstruction of intact human integrin alphaIIbbeta3: new implications for activation-dependent ligand binding. Blood 122, 4165–4171, doi:10.1182/blood-2013-04-499194 (2013).

19 Dhankhar, P., Trinh, T. K. H., Qiu, W. & Guo, Y. Characterization of Ca(2+)-ATPase, LMCA1, with native cell membrane nanoparticles system. Biochim Biophys Acta Biomembr 1865, 184143, doi:10.1016/j.bbamem.2023.184143 (2023).

20 Adair, B. D., Xiong, J. P., Yeager, M. & Arnaout, M. A. Cryo-EM structures of full-length integrin alphaIIbbeta3 in native lipids. Nat Commun 14, 4168, doi:10.1038/s41467-023-39763-0 (2023).

21 Sivasakthi, V. et al. Aromatic-aromatic interactions: analysis of pi-pi interactions in interleukins and TNF proteins. Bioinformation 9, 432–439, doi:10.6026/97320630009432 (2013).

22 Gallivan, J. P. & Dougherty, D. A. Cation-pi interactions in structural biology. Proc Natl Acad Sci U S A 96, 9459–9464, doi:10.1073/pnas.96.17.9459 (1999).

23 Arnaout, M. A. Integrins: A Bedside to Bench to Bedside Story. Trans Am Clin Climatol Assoc 133, 34–55 (2023).

24 Cody, V. et al. Structural Analysis of the Complex of Human Transthyretin with 3’,5’-Dichlorophenylanthranilic Acid at 1.5 A Resolution. Molecules 27, doi:10.3390/molecules27217206 (2022).

25 Vickers, S. et al. In vitro and in vivo studies on the metabolism of tirofiban. Drug Metab Dispos 27, 1360–1366 (1999).

26 Peerlinck, K. et al. MK-383 (L-700,462), a selective nonpeptide platelet glycoprotein IIb/IIIa antagonist, is active in man. Circulation 88, 1512–1517, doi:10.1161/01.cir.88.4.1512 (1993).

27 Hantgan, R. R., Stahle, M. C. & Lord, S. T. Dynamic regulation of fibrinogen: integrin alphaIIbbeta3 binding. Biochemistry 49, 9217–9225, doi:10.1021/bi1009858 (2010).

28 Scarborough, R. M. & Gretler, D. D. Platelet glycoprotein IIb-IIIa antagonists as prototypical integrin blockers: novel parenteral and potential oral antithrombotic agents. J Med Chem 43, 3453–3473, doi:10.1021/jm000022w (2000).

29 Mousa, S. A. et al. Antiplatelet efficacy of XV459, a novel nonpeptide platelet GPIIb/IIIa antagonist: comparative platelet binding profiles with c7E3. J Pharmacol Exp Ther 286, 1277–1284 (1998).

30 Litvinov, R. I., Farrell, D. H., Weisel, J. W. & Bennett, J. S. The Platelet Integrin alphaIIbbeta3 Differentially Interacts with Fibrin Versus Fibrinogen. J Biol Chem 291, 7858–7867, doi:10.1074/jbc.M115.706861 (2016).

31 Welsh, J. D. et al. Hemodynamic regulation of perivalvular endothelial gene expression prevents deep venous thrombosis. J Clin Invest 129, 5489–5500, doi:10.1172/JCI124791 (2019).

32 Hoofnagle, M. H. et al. Defects in vein valve PROX1/FOXC2 antithrombotic pathway in endothelial cells drive the hypercoagulable state induced by trauma and critical illness. J Trauma Acute Care Surg 95, 197–204, doi:10.1097/TA.0000000000003945 (2023).

33 Diaz, J. A. et al. Choosing a mouse model of venous thrombosis: a consensus assessment of utility and application. J Thromb Haemost 17, 699–707, doi:10.1111/jth.14413 (2019).

34 Tutwiler, V. et al. Kinetics and mechanics of clot contraction are governed by the molecular and cellular composition of the blood. Blood 127, 149–159, doi:10.1182/blood-2015-05-647560 (2016).

35 Tutwiler, V., Wang, H., Litvinov, R. I., Weisel, J. W. & Shenoy, V. B. Interplay of Platelet Contractility and Elasticity of Fibrin/Erythrocytes in Blood Clot Retraction. Biophys J 112, 714–723, doi:10.1016/j.bpj.2017.01.005 (2017).

36 Leon, C. et al. Megakaryocyte-restricted MYH9 inactivation dramatically affects hemostasis while preserving platelet aggregation and secretion. Blood 110, 3183–3191, doi:10.1182/blood-2007-03-080184 (2007).

37 Cohen, I., Burk, D. L. & White, J. G. The effect of peptides and monoclonal antibodies that bind to platelet glycoprotein IIb-IIIa complex on the development of clot tension. Blood 73, 1880–1887 (1989).

38 Osdoit, S. & Rosa, J. P. Fibrin clot retraction by human platelets correlates with alpha(IIb)beta(3) integrin-dependent protein tyrosine dephosphorylation. J Biol Chem 276, 6703–6710, doi:10.1074/jbc.M008945200 (2001).

39 Lin, F. Y. et al. A general chemical principle for creating closure-stabilizing integrin inhibitors. Cell 185, 3533–3550 e3527, doi:10.1016/j.cell.2022.08.008 (2022).

40 Benet, L. Z. & Hoener, B. A. Changes in plasma protein binding have little clinical relevance. Clin Pharmacol Ther 71, 115–121, doi:10.1067/mcp.2002.121829 (2002).

41 Smith, D. A., Di, L. & Kerns, E. H. The effect of plasma protein binding on in vivo efficacy: misconceptions in drug discovery. Nat Rev Drug Discov 9, 929–939, doi:10.1038/nrd3287 (2010).

42 Liu, X., Wright, M. & Hop, C. E. Rational use of plasma protein and tissue binding data in drug design. J Med Chem 57, 8238–8248, doi:10.1021/jm5007935 (2014).

43 Di, L. An update on the importance of plasma protein binding in drug discovery and development. Expert Opin Drug Discov 16, 1453–1465, doi:10.1080/17460441.2021.1961741 (2021).

44 Yan, Z. et al. New Methodology for Determining Plasma Protein Binding Kinetics Using an Enzyme Reporter Assay Coupling with High-Resolution Mass Spectrometry. Anal Chem 95, 4086–4094, doi:10.1021/acs.analchem.2c04864 (2023).

45 Schwarz, M. et al. Conformation-specific blockade of the integrin GPIIb/IIIa: a novel antiplatelet strategy that selectively targets activated platelets. Circ Res 99, 25–33, doi:10.1161/01.RES.0000232317.84122.0c (2006).

46 Ndrepepa, G. et al. Correlates of poor outcome among patients with bleeding after coronary interventions. Coron Artery Dis 25, 456–462, doi:10.1097/MCA.0000000000000126 (2014).

47 Baba, K., Aga, Y., Nakanishi, T., Motoyama, T. & Ueno, H. UR-3216: a manageable oral GPIIb/IIIa antagonist. Cardiovasc Drug Rev 19, 25–40, doi:10.1111/j.1527-3466.2001.tb00181.x (2001).

48 Michel, M. C., Michel-Reher, M. B. & Hein, P. A Systematic Review of Inverse Agonism at Adrenoceptor Subtypes. Cells 9, doi:10.3390/cells9091923 (2020).

49 Meye, F. J., Trezza, V., Vanderschuren, L. J., Ramakers, G. M. & Adan, R. A. Neutral antagonism at the cannabinoid 1 receptor: a safer treatment for obesity. Mol Psychiatry 18, 1294–1301, doi:10.1038/mp.2012.145 (2013).

50 Seiffert, D. et al. Regulation of clot retraction by glycoprotein IIb/IIIa antagonists. Thromb Res 108, 181–189, doi:10.1016/s0049-3848(02)00395-x (2002).

51 Caen, J. P. et al. Congenital bleeding disorders with long bleeding time and normal platelet count: I. Glanzmann’s thrombasthenia (report of fifteen patients). The American Journal of Medicine 41, 4–26 (1966).

52 Law, D. A. et al. Integrin cytoplasmic tyrosine motif is required for outside-in alphaIIbbeta3 signalling and platelet function. Nature 401, 808–811, doi:10.1038/44599 (1999).

53 Jiang, J., Wang, W., Sane, D. C. & Wang, B. Synthesis of RGD analogs as potential vectors for targeted drug delivery. Bioorg Chem 29, 357–379, doi:10.1006/bioo.2001.1227 (2001).

54 Punjani, A., Rubinstein, J. L., Fleet, D. J. & Brubaker, M. A. cryoSPARC: algorithms for rapid unsupervised cryo-EM structure determination. Nat Methods 14, 290–296, doi:10.1038/nmeth.4169 (2017).

55 Afonine, P. V. et al. phenix.model_vs_data: a high-level tool for the calculation of crystallographic model and data statistics. J Appl Crystallogr 43, 669–676, doi:10.1107/S0021889810015608 (2010).

56 Emsley, P., Lohkamp, B., Scott, W. G. & Cowtan, K. Features and development of Coot. Acta Crystallogr D Biol Crystallogr 66, 486–501, doi:10.1107/S0907444910007493 (2010).

57 Mussbacher, M. et al. Optimized plasma preparation is essential to monitor platelet-stored molecules in humans. PLoS One 12, e0188921, doi:10.1371/journal.pone.0188921 (2017).

58 Bennett, J. S. & Vilaire, G. Exposure of platelet fibrinogen receptors by ADP and epinephrine. J Clin Invest 64, 1393–1401, doi:10.1172/JCI109597 (1979).

59 Tucker, K. L., Sage, T. & Gibbins, J. M. Clot retraction. Methods Mol Biol 788, 101–107, doi:10.1007/978-1-61779-307-3_8 (2012).

60 Gapizov, S. S. et al. Fusion with an albumin-binding domain improves pharmacokinetics of an alphavbeta3-integrin binding fibronectin scaffold protein. Biotechnol Appl Biochem 66, 617–625, doi:10.1002/bab.1762 (2019).

61 Gollomp, K. et al. Neutrophil accumulation and NET release contribute to thrombosis in HIT. JCI Insight 3, doi:10.1172/jci.insight.99445 (2018).

62 Hui, K. Y., Haber, E. & Matsueda, G. R. Monoclonal antibodies to a synthetic fibrin-like peptide bind to human fibrin but not fibrinogen. Science 222, 1129–1132, doi:10.1126/science.6648524 (1983).

